# A Myosin Nanomotor Essential for Stereocilia Maintenance Expands the Etiology of Hereditary Hearing Loss DFNB3

**DOI:** 10.1101/2025.02.19.639121

**Authors:** Ghazaleh Behnammanesh, Abigail K. Dragich, Xiayi Liao, Shadan Hadi, Mi-Jung Kim, Benjamin Perrin, Shinichi Someya, Gregory I. Frolenkov, Jonathan E. Bird

## Abstract

Cochlear hair cells transduce sound using stereocilia, and disruption to these delicate mechanosensors is a significant cause of hearing loss. Stereocilia architecture is dependent upon the nanomotor myosin 15. A short isoform (MYO15A-2) drives stereocilia development by delivering an elongation-promoting complex (EC) to stereocilia tips, and an alternatively spliced long isoform (MYO15A-1) tunes postnatal size in shorter stereocilia, which possess mechanosensitive ion channels. Disruption of these functions causes two distinct stereocilia pathologies, which underly human autosomal recessive non-syndromic hearing loss DFNB3. Here, we characterize a new isoform, MYO15A-3, that increases expression in postnatal hair cells as the developmental MYO15A-2 isoform wanes reciprocally. We show the critical EC complex is initially delivered by MYO15A-2, followed by a postnatal handover to MYO15A-3, which continues to deliver the EC. In a *Myo15a-3* mutant mouse, stereocilia develop normally with correct EC targeting, but lack the EC postnatally and do not maintain their adult architecture, leading to progressive hearing loss. We conclude MYO15A-3 delivers the EC in postnatal hair cells and that the EC and MYO15A-3 are both required to maintain stereocilia integrity. Our results add to the spectrum of stereocilia pathology underlying DFNB3 hearing loss and reveal new molecular mechanisms necessary for resilient hearing during adult life.

## INTRODUCTION

Myosin 15 (MYO15A) is a cytoskeletal nanomotor that is expressed by inner ear hair cells is required for the assembly of mechanosensitive stereocilia, which detect sound and accelerations of the head (Moreland and Bird 2022). In humans, *MYO15A* mutations cause autosomal recessive non-syndromic deafness DFNB3, which is characterized by congenital, profound hearing loss with varying degrees of vestibular dysfunction (Winata et al. 1995; Friedman et al. 1995; A. Wang et al. 1998). Extensive research into the molecular mechanisms of DFNB3 hearing loss reveal a central role for MYO15A in regulating stereocilia growth. Each cochlear hair cell arranges approximately 100 actin-based stereocilia into three rows of ascending height to form a functional mechanosensory hair bundle (Caprara and Peng 2022). Stereocilia emerge from hair cell microvilli and must lengthen and widen to develop into their mature form (Krey et al. 2023; Lewis G. Tilney et al. 1986; Kaltenbach, Falzarano, and Simpson 1994; Zine and Romand 1996; Krey et al. 2020). A *Myo15a* missense substitution in the *shaker-2 (sh2)* mouse prevents normal stereocilia growth and has provided the prototypical model to explain DFNB3 pathology (Probst et al. 1998). Clinically, several hundred mutations in *MYO15A* are reported to cause DFNB3 (Rehman et al. 2016; J. Zhang et al. 2019; Miyoshi et al. 2024).

The mechanism of stereocilia growth involves trafficking of MYO15A along actin filaments (Belyantseva, Boger, and Friedman 2003; Bird et al. 2014; Jiang et al. 2021) and its accumulation at the distal tips of nascent stereocilia (Belyantseva, Boger, and Friedman 2003; Belyantseva et al. 2005), a major site of actin polymerization (Schneider et al. 2002; Rzadzinska et al. 2004; Drummond et al. 2015; McGrath et al. 2021). MYO15A delivers a complex (EC) of epidermal growth factor receptor pathway substrate 8 (EPS8), whirlin (WHRN), G protein subunit alpha i3 (GNAI3), and G-protein signaling modulator 2 (GPSM2), termed the elongation complex, which establishes the molecular identity and differential elongation of the tallest row 1 stereocilia (Delprat et al. 2005; Belyantseva et al. 2005; Manor et al. 2011; Mauriac et al. 2017; Tadenev et al. 2019). Although MYO15A and the EC proteins are each individually required for stereocilia elongation; the exact mechanisms regulating this actin polymerization are not well understood. EPS8 can influence actin filament elongation by binding actin filament barbed ends and acting as a leaky capping protein, which allows new monomer addition (Disanza et al. 2004; Gaeta et al. 2021). The ATPase domain of MYO15A can also directly catalyze the nucleation of new actin filaments that contribute to stereocilia elongation (Gong et al. 2022; Moreland et al. 2025). The relative balance of these actin-regulatory activities and how they are integrated at the stereocilia tip is still unclear.

The *MYO15A* locus contains 66 exons; alternative splicing of the 3.6 kb exon 2 leads to the translation of two distinct protein isoforms; MYO15A-1 (a.k.a. MYO15A-L), with a large 133 kDa N-terminal domain, and MYO15A-2 (a.k.a. MYO15A-S) (Fig. 1A) (Liang et al. 1999; Belyantseva, Boger, and Friedman 2003; Fang et al. 2015). Both isoforms share common structural motifs including an actin-activated ATPase domain, a tail domain containing light chain binding sites, two myosin tail homology 4 (MyTH4) domains, a single Src-homology 3 (SH3) domain and two protein 4.1, ezrin, radixin and moesin (FERM) domains (Fig. 1B). These two isoforms have independent functions that were originally masked by the stunted stereocilia phenotype of *Myo15a^sh2/sh2^* mice, which was caused by a mutation in the shared ATPase domain that inactivates both MYO15A-1 and MYO15A-2 (Probst et al. 1998; Belyantseva, Boger, and Friedman 2003; Fang et al. 2015). An isoform-specific deletion of MYO15A-1 in the *Myo15a*^Δ*N*^ mouse showed the largest isoform is dispensable for the initial development of graded stereocilia heights and mechanotransduction (MET) currents (Fang et al. 2015). These results suggest that remaining MYO15A-2 was sufficient to drive initial stereocilia elongation (Fang et al. 2015). Consistent with this observation, EPS8 and WHRN were correctly localized in *Myo15a^(^*^Δ*N/*Δ*N)*^ hair cells (Fang et al. 2015). Although MYO15A-1 is dispensable for development, it is required for maintaining the adult architecture of the shorter rows 2 and 3 transducing stereocilia with active MET channels (Fang et al. 2015). This function reveals a second distinct DFNB3 pathology, which is caused by variants specifically targeting the large *MYO15A* exon 2 (Nal et al. 2007).

**Figure 1.**
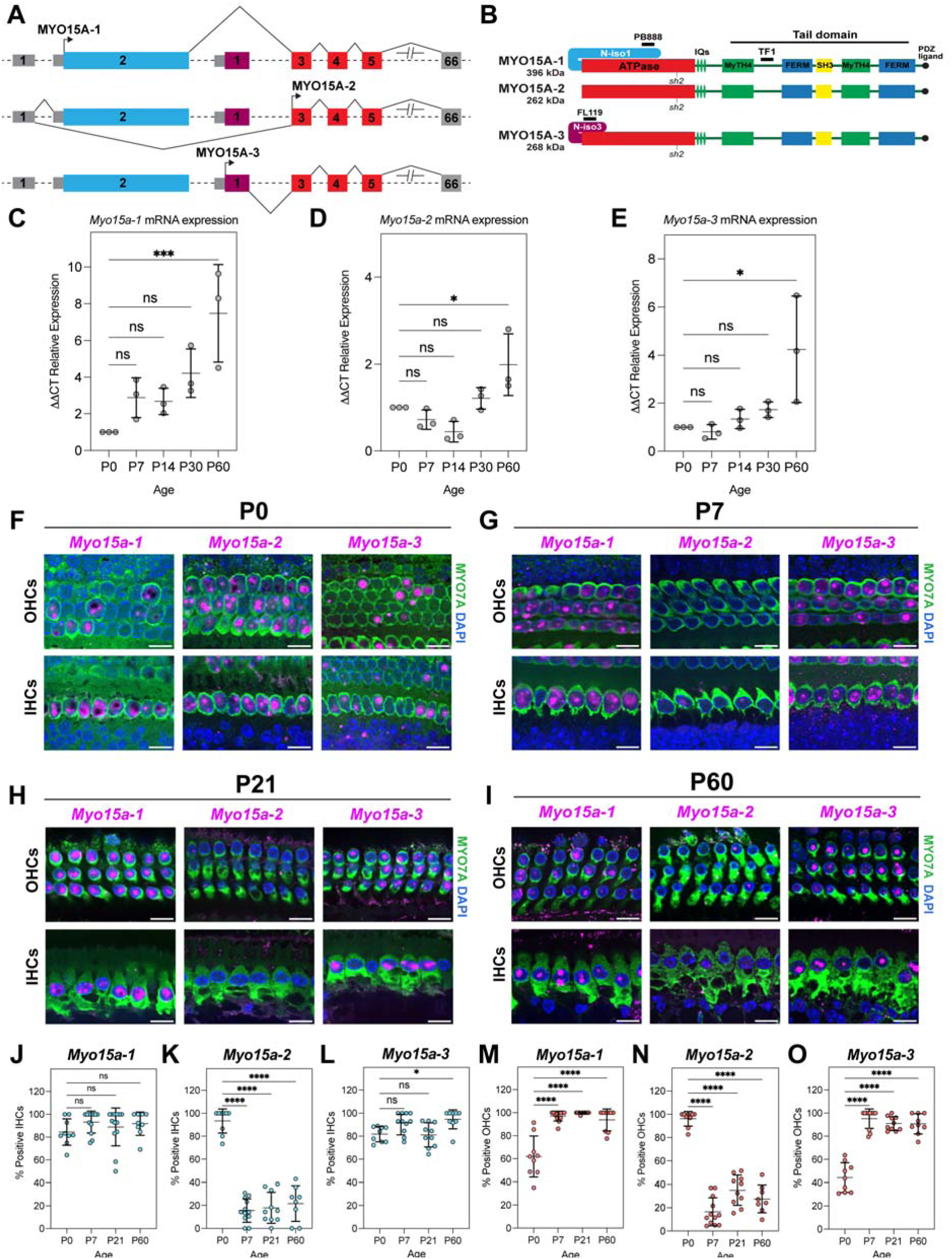
Expression of *Myo15a* isoforms in mouse cochleae. **(A)** Cartoon representation of the *Myo15a* gene structure with alternative transcription start sites. **(B)** MYO15A isoforms share identical ATPase and tail domains but have unique N-terminal domains. **(C-E)** Quantitative RT-PCR analysis of *Myo15*a mRNA at key stages of hair bundle development and maturation in wild-type C57BL/6J mouse cochleae. Expression of *Myo15a* transcripts are normalized to *Hprt* and P0 (ΔΔC_T_). Data points are mean ± SD (n = 3 independent animals per group). **(F-I)** BaseScope *in situ* hybridization of *Myo15a* isoforms in P0, P7, P21, and P60 mouse cochlea (magenta). IHCs and OHCs are identified with anti-MYO7A labeling (green), and nuclei are stained with DAPI (blue). **(J-O)** Quantification of BaseScope signal in IHCs **(J, K, L)** and OHCs (**M, N, O**). Each data point is one measurement, mean ± SD (n = 3 independent animals per group). Statistical significance was determined by one-way ANOVA with Dunnett’s multiple comparison test, **** P<0.0001, ***< 0.001, *P<0.05. Scale bars: 10 µm.

Further interpretation of these results is complicated by evidence that hair cells express an additional MYO15A isoform. Using 5’-RACE, we previously identified an unannotated exon (mm10 chr11:60,480,418–60,480,621) and transcription start site (TSS) in mouse and human pituitary RNA encoding a distinct 6 kDa N-terminal extension to the ATPase domain (Rehman et al. 2016). The full-length *Myo15a-3* transcript was further detected in cochlear RNA (Abu Rayyan et al. 2020) and in single-cell nanopore RNAseq of hair cells (Ranum et al. 2019). Although the function of MYO15A-3 in the cochlea is unknown; a deeply intronic *MYO15A* variant predicted to abolish splice donor activity of *MYO15A-3* exon 1 was identified in multiple pedigrees with DFNB3-linked hearing loss, suggesting it is required for hearing (Abu Rayyan et al. 2020).

To explore the potential involvement of MYO15A-3 in DFNB3 pathology, we generated a mutant mouse with a specific deletion of *Myo15a-3,* leaving the expression of *Myo15a-1* and *Myo15a-2* intact. *Myo15a-3* knock-out mice initially developed normal stereocilia hair bundles but exhibited progressive hearing loss. Our experiments revealed that EC proteins are dependent upon two MYO15A isoforms for their correct stereocilia targeting; EC proteins engage MYO15A-2 in neonatal hair cells, then transitioned to MYO15A-3 use postnatally. In the absence of this EC handover from MYO12A-2 to MYO15A-3, however, there was selective degeneration of the tallest row 1 stereocilia. This demonstrates that MYO15A-3 and the EC are necessary for continued maintenance of the mature hair bundle architecture maintenance. Our results reveal how three MYO15A isoforms cooperate to build and sustain hair cell mechanosensation, and expand the spectrum of pathologies that underlie the human deafness DFNB3.

## RESULTS

### *Myo15a-3* is expressed postnatally by cochlear hair cells

To explore the expression profile of *Myo15a-3*, we analyzed the relative abundance of *Myo15a* mRNA transcripts using qPCR in wild-type (WT) C57BL/6J mouse cochlear RNA at key developmental stages: P0, P7, P14, P30, and P60. Previous studies have shown developmental regulation of *Myo15a* isoform transcription; *Myo15a-2* is expressed embryonically and declines after P0 and *Myo15a-1* expressed postnatally from P4.5 onwards (Fang et al. 2015; Krey et al. 2020). We designed and validated primers to uniquely detect *Myo15a-1, -2* and *-3* using SYBR Green chemistry (Fig. S1). Using this approach, we detected a steady increase in *Myo15a-3* transcript abundance relative to P0, with a statistically significant 5-fold increase at P60 (Fig. 1E). Transcripts for *Myo15a-1* followed a similar postnatal time-course with a statistically significant 8-fold increase detected by P60 (Fig. 1C). In contrast, *Myo15a-2* transcripts were decreased by P7 and P14, although this trend was not statistically significant (Fig. 1D). Unexpectedly, *Myo15a-2* transcripts were significantly increased 2-fold by P60 (Fig. 1D).

To determine the identity of cells expressing *Myo15a-3* in the cochlea, we performed BaseScope *in situ* hybridization in C57BL/6J wild-type mice at P0 (Fig. 1F), P7 (Fig. 1G), P21 (Fig. 1H), and P60 (Fig. 1I). BaseScope probes were designed to anneal to exon-exon boundaries and uniquely discriminate between the *Myo15a-1, -2* and *-3* transcripts (Fig. S1). To ensure the specificity and accuracy of hybridization, BaseScope signals were compared to mouse peptidylprolyl isomerase B (*Ppib*) and a bacterial dihydrodipicolinate reductase (*dapB*) as positive and negative controls, respectively (Fig. S1). *Myo15a-3* mRNA transcripts were detected as early as P0 in MYO7A-positive inner hair cells (IHCs) and outer hair cells (OHCs) (Figs. 1F, L, O). By P7, the majority of IHCs and OHCs were positive for *Myo15a-3* (Figs. 1G, L, O), and continued to express *Myo15a-3* at P21 (Fig. 1H) and P60 (Fig. 1I). *Myo15a-1* was expressed in a similar spatial-temporal pattern, with initial detection in IHCs and OHCs at P0 (Fig. 1F, J, M), expanding to the majority of IHCs and OHCs by P7 (Fig. 1G), and persisting up until at least P60 (Fig. 1I). Conversely, *Myo15a-2* expression was detected in almost all IHCs and OHCs at P0 (Fig. 1F, K, N). Despite the overall number of IHCs and OHCs expressing *Myo15a-2* being significantly reduced by P7, a persistent fraction of OHCs and IHCs were still positive from P7 – P60 (Fig. 1G, H, I). No *Myo15a* transcripts were detected in any other cell type in the organ of Corti. We conclude that *Myo15a-3* is specifically expressed in hair cells, and that in concert with *Myo15a-1*, is continually expressed into adulthood.

### *Myo15a-3* null mice have a rapidly progressive hearing loss

Existing models of DFNB3 human deafness utilize mice with nonsense substitutions specific to *Myo15a-1* (p.E1086*, Δ*N*), or missense substitutions/deletions that target exons common to *Myo15a-1*, *Myo15a-2* and *Myo15a-3* such as the *shaker-2* (p.C1779Y, *sh2*), *jordan* (p.D1647G, *jd*) and *shaker-2^J^* (*sh2-J*) mutations (Anderson et al. 2000; Fang et al. 2015; Probst et al. 1998; Belyantseva et al. 2003; Stepanyan et al. 2006; Moreland et al. 2025). Based upon its reported exon usage (Ranum et al. 2019; Rehman et al. 2016), the function of *Myo15a-3* is also expected to be impacted by the *sh2, jd*, and *sh2-J* mutant alleles. To specifically study the function of *Myo15a-3* in the auditory system, we used the CRISPR/Cas9 system to delete exon 1 containing the transcription start site (TSS) unique to *Myo15a-3* by non-homologous end joining (Fig. S2). No other known *Myo15a* transcript utilizes this exon, and its deletion was predicted to specifically target *Myo15a-3*. Additionally, an intronic DFNB3 variant was computationally predicted to ablate the splice donor activity of exon 1 (Abu Rayyan et al. 2020). Following germline transmission, Sanger sequencing of a genomic PCR amplicon spanning the targeted genomic region confirmed a 350 bp deletion that eliminated exon 1 of *Myo15a-3.* We call this allele *Myo15a^(^*^Δ*3)*^. Male and female *Myo15a^(^*^Δ*3/*Δ*3)*^ mice were healthy, fertile, and born in normal Mendelian ratios.

To test if *Myo15a-3* is required for hearing function, we performed longitudinal auditory brainstem response (ABR) measurements in wild-type *Myo15a^(+/+)^*, heterozygous *Myo15a^(+/^*^Δ*3)*^, and homozygous *Myo15a^(^*^Δ*3/*Δ*3)*^ littermates at P17 (Fig. 2A), P30 (Fig. 2B), and P60 (Fig. 2C). At P17, the earliest age where hearing thresholds can be reproducibly measured, we detected a small but statistically significant elevated threshold to broadband click stimuli in the *Myo15a^(^*^Δ*3/*Δ*3)*^ cohort (46 ± 12 dB SPL, mean ± S.D), compared to wild-type *Myo15a^(+/+)^* littermate controls (35 ± 5 dB SPL). More detailed analysis using pure-tone stimuli at P17 revealed the broadband click threshold shift in *Myo15a^(^*^Δ*3/*Δ*3)*^ mice likely originated from a large threshold shift at 32 kHz (88 ± 5 dB SPL) versus wild-type *Myo15a^(+/+)^* littermate controls (35 ± 9 dB SPL), plus additional small but statistically significant threshold elevations at 8 and 16 kHz (Fig. 2A, S3A). No statistically significant differences in threshold shifts were observed between *Myo15a^(+/+)^*and *Myo15a^(+/^*^Δ*3)*^ cohorts for click or pure-tone thresholds at any age (Fig. 2A, B, C). Broadband click and pure-tone thresholds for *Myo15a^(^*^Δ*3/*Δ*3)*^ mice were all elevated close to the instrument’s maximum sound pressure (90 dB SPL) by P30 and P60, indicating severe hearing loss across all frequencies (Fig. 2B, C, S3B, C). At P30 and P60, elevated ABR thresholds were also detected at 32 kHz in *Myo15a^(+/+)^* mice compared to *Myo15a^(+/^*^Δ*3)*^ littermates (Fig. 2B, C). We observed no effect of biological sex upon hearing loss in *Myo15a^(^*^Δ*3/*Δ*3)*^ mice at any age, by pure-tone stimulus (Fig. S3D) or broadband click (Fig. S3E).

**Figure 2.**
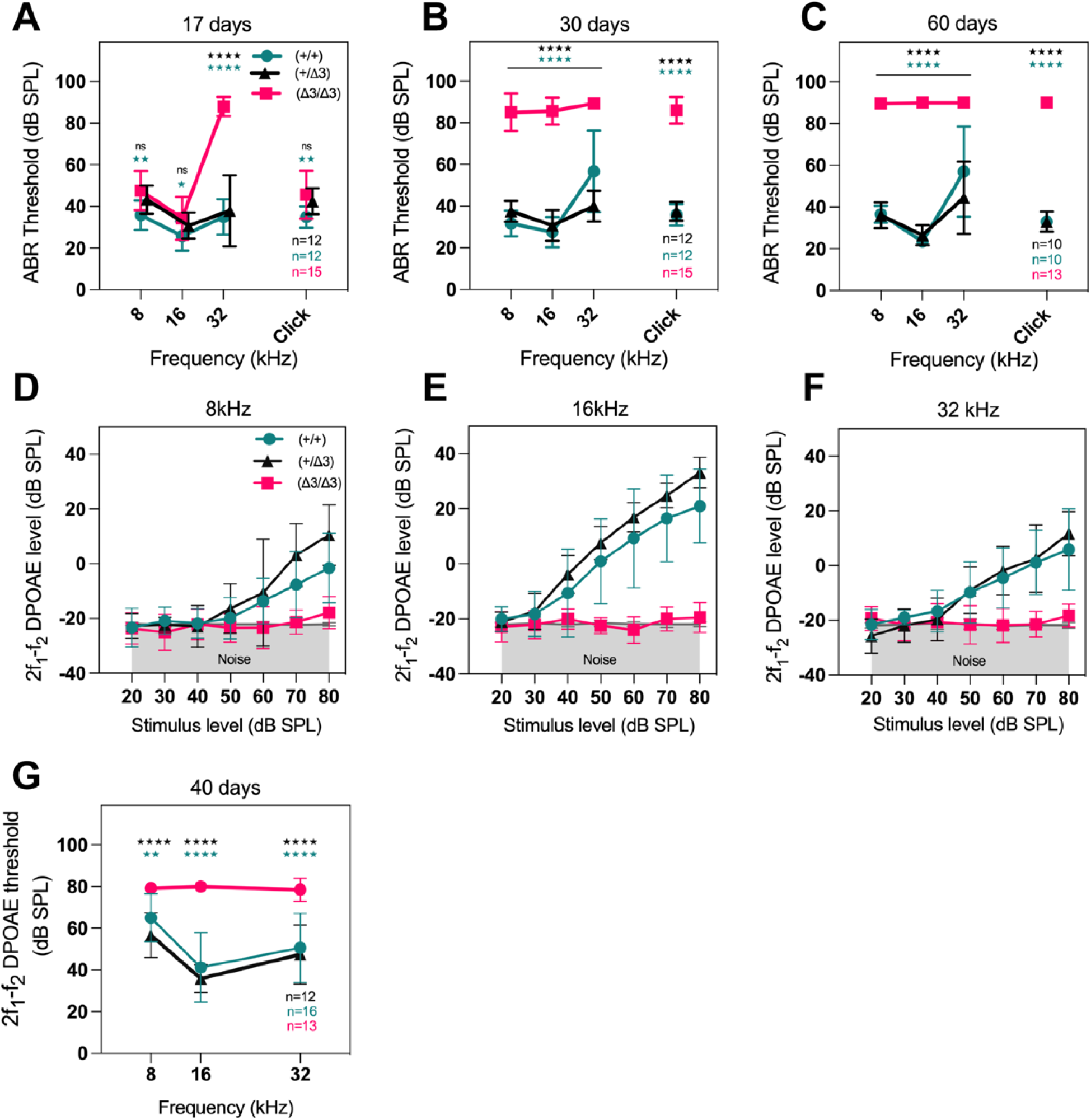
Auditory testing in mutant *Myo15a-3* null mice. **(A-C)** Longitudinal auditory brainstem response (ABR) thresholds in *Myo15a-3* littermates at 17 **(A)**, 30 **(B)** and 60 days **(C)**. Statistical comparisons are indicated by green asterisks (*Myo15a^(+/+)^* vs *Myo15a^(^*^Δ*3/*Δ*3)*^) and black asterisks (*Myo15a^(+/^*^Δ*3)*^ vs *Myo15a^(^*^Δ*3/*Δ*3)*^). **(D-F)** Distortion product otoacoustic emission (DPOAE) amplitude and thresholds were measured at 8 **(D)**, 16 **(E)**, and 32 kHz **(F)** in *Myo15a^(+/+)^* n=16 (8 female and 8 male), *Myo15a^(+/^*^Δ*3*)^ n=12 (8 female and 4 male), and *Myo15a^(^*^Δ*3/*Δ*3)*^ n=13 (6 female and 7 male) mice at P40. *Myo15a^(^*^Δ*3/*Δ*3)*^ mice demonstrated significant DPOAE threshold increase across all tested frequencies. All data are presented as mean ± SD, and statistical analyses were conducted using two-way ANOVA with Tukey’s multiple comparisons test. Significance at all frequencies is indicated by ** P < 0.01 and **** P < 0.0001. As with the ABR thresholds, the color-coding of asterisks (black for *Myo15a^(^*^Δ*3/*Δ*3)*^ vs. *Myo15a^(+/+)^*, and green for *Myo15a^(^*^Δ*3/*Δ*3*)^ vs. *Myo15a^(+/^*^Δ*3*)^) signifies the specific genotype comparisons for P-value.

As an additional measure of hearing function, distortion product otoacoustic emissions (DPOAEs) were recorded to examine active cochlear amplification in *Myo15a^(+/+)^*, *Myo15a^(+/^*^Δ*3)*^ and *Myo15a^(^*^Δ*3/*Δ*3)*^ littermates at P40 (Fig. S3F). DPOAEs originate from the nonlinear amplification of sound by OHCs and are generated in response to two primary tones (f1, f2) presented simultaneously. DPOAE amplitudes (2f1 - f2) were absent at all frequencies tested (8, 16, and 32 kHz) in *Myo15a^(^*^Δ*3/*Δ*3)*^ mice compared with *Myo15a^(+/^*^Δ*3)*^ and *Myo15a^(+/+)^* littermates (Fig. 2D-F), indicating a significant impairment of OHC function (Fig. 2G). In summary, our results show that the *Myo15a^(^*^Δ*3)*^ allele causes a recessive hearing loss with a moderate ABR phenotype at P17, that rapidly progresses to severe hearing loss by P30. We conclude that *Myo15a-3* is not required for the initial acquisition of hearing function but is instead necessary for its maturation and maintenance.

### Disruption of *Myo15a-3* does not interfere with the expression of other *Myo15a* isoforms

We deleted exon 1 containing the TSS of *Myo15a-3* to specifically ablate this isoform (Fig. S2). To validate that our targeting strategy was specific to *Myo15a-3*, we performed BaseScope *in situ* hybridization at key developmental stages to detect *Myo15a-1*, *Myo15a-2* and *Myo15a-3* in mutant *Myo15a^(^*^Δ*3/*Δ*3)*^ cochleae and normal hearing *Myo15a^(+/^*^Δ*3)*^ littermate controls. We assayed *Myo15a-2* at P0, the peak time of expression in WT hair cells (Fig. 3A) and found no statistically significant difference in the percentage of *Myo15a-2*-positive hair cells in *Myo15a^(^*^Δ*3/*Δ*3)*^ versus *Myo15a^(+/^*^Δ*3)*^ controls (Fig. 3D). We assayed *Myo15a-1* at P21, when expression of this isoform was high in WT hair cells (Fig. 3B) and again saw no statistically significant difference in the percentage of positive hair cells (Fig. 3E). Finally, we measured *Myo15a-3* at P21, when this transcript is abundantly expressed in WT hair cells and observed a total loss of BaseScope signal in both IHCs and OHCs of *Myo15a^(^*^Δ*3/*Δ*3)*^ cochlea versus *Myo15a^(+/^*^Δ*3)*^ controls (Fig. 3C, F). These data confirm the specificity of our targeting approach and that *Myo15a^(^*^Δ*3/*Δ*3)*^ hair cells do not express *Myo15a-3*. Our results further show that the hearing loss observed in *Myo15a^(^*^Δ*3/*Δ*3)*^ mice is not due to a general disruption of the *Myo15a* transcriptional landscape but specifically caused by the loss of *Myo15a-3*.

**Figure 3.**
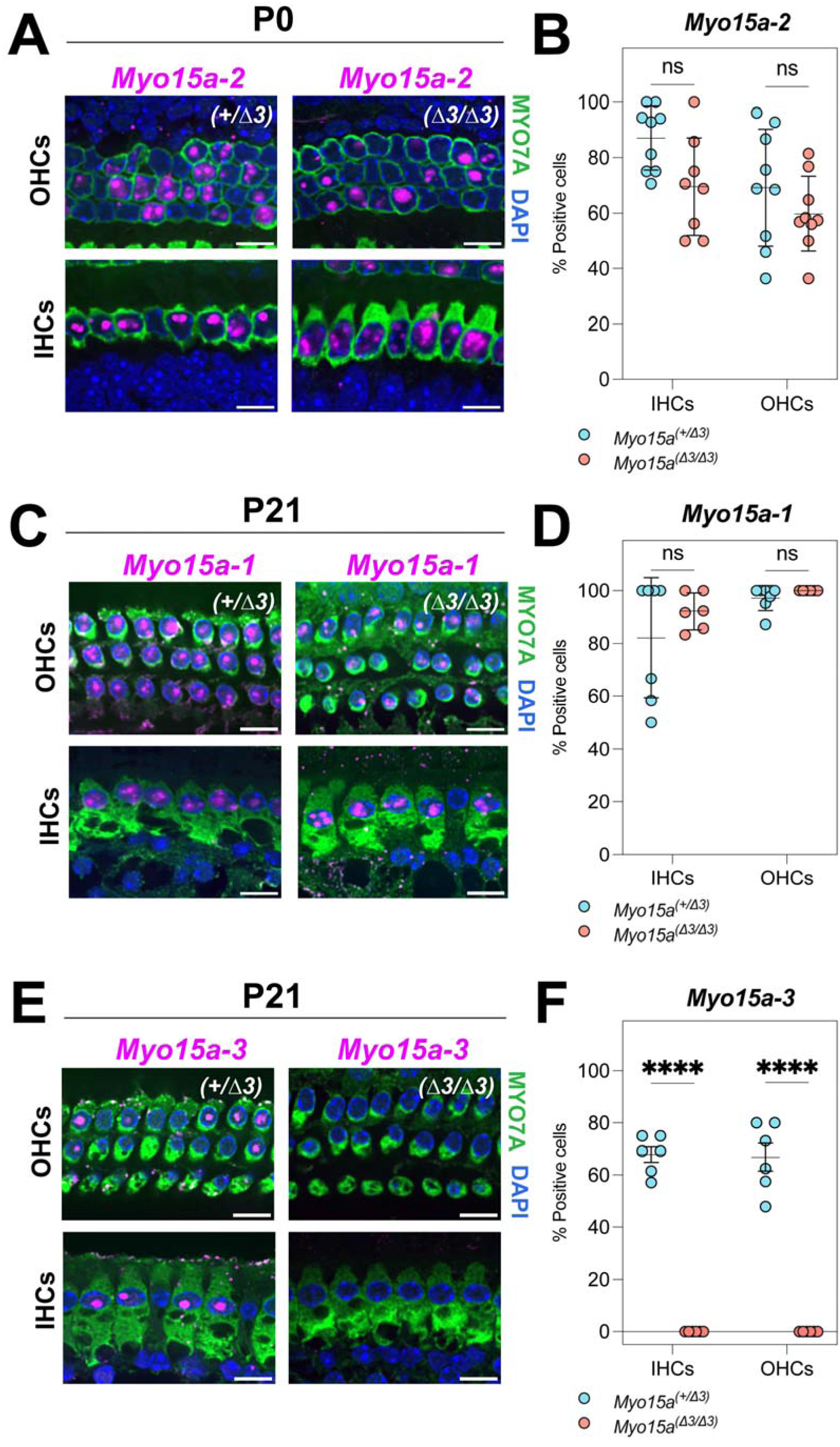
The deletion of *Myo15a-3* does not affect the expression of other *Myo15a* isoforms. **(A, B)** The BaseScope signal shows the expression of *Myo15a-1* and *Myo15a-2* in IHCs and OHCs of *Myo15a^(+/^*^Δ*3)*^ and *Myo15a^(^*^Δ*3/*Δ*3)*^ mice at P0 and P21. **(C)** The expression of *Myo15a-3* in IHCs and OHCs of P21 *Myo15a^(+/^*^Δ*3)*^. Deletion of exon 1 of *Myo15a-3* results in the complete loss of *Myo15a-3* expression in *Myo15a^(^*^Δ*3/*Δ*3)*^ mice. **(D-F)** The comparison of the percentage of IHCs and OHCs expressing *Myo15a* isoforms between *Myo15a^(+/^*^Δ*3)*^ and *Myo15a^(^*^Δ*3/*Δ*3)*^ mice. Quantitative data are presented as mean ± SD from n=2-3 mice (males and females) per genotype. Statistical significance was determined by two-way ANOVA with Šidák and Fisher’s LSD multiple comparison tests, **** P<0.0001. Scale bars: 10 µm.

### MYO15A-3 is targeted to the tallest row 1 stereocilia

Having confirmed that *Myo15a-3* is required for hearing, we next studied the subcellular localization of MYO15A-3 protein to understand its function within cochlear hair cells. Other MYO15A isoforms have previously been reported to be concentrated at the distal tips of hair cell stereocilia, including MYO15A-1 (Fang et al. 2015; Krey et al. 2020) and MYO15A-2 (Belyantseva et al. 2005; Fang et al. 2015; Belyantseva, Boger, and Friedman 2003; Rzadzinska et al. 2004). To detect MYO15A-3 in hair cells, we cultured C57BL/6J wild-type cochleae at P5 as organotypic explants and used injectoporation (Xiong et al. 2014) to overexpress plasmid DNA encoding EGFP-tagged MYO15A-3. Following injectoporation, we observed EGFP-MYO15A-3 fluorescence strongly concentrated at the tips of the tallest row 1 stereocilia in IHCs and OHCs (Fig. 4A). Less intense EGFP-MYO15A-3 signal was observed at the tips of the shorter stereocilia rows and 2 and 3 and in hair cell microvilli (Fig. 4A). We conclude that MYO15A-3 primarily localizes to the tip compartment of row 1 stereocilia in hair cells, which is distinct from MYO15A-1, which concentrates in the shorter rows 2 and 3 (Fang et al. 2015).

**Figure 4.**
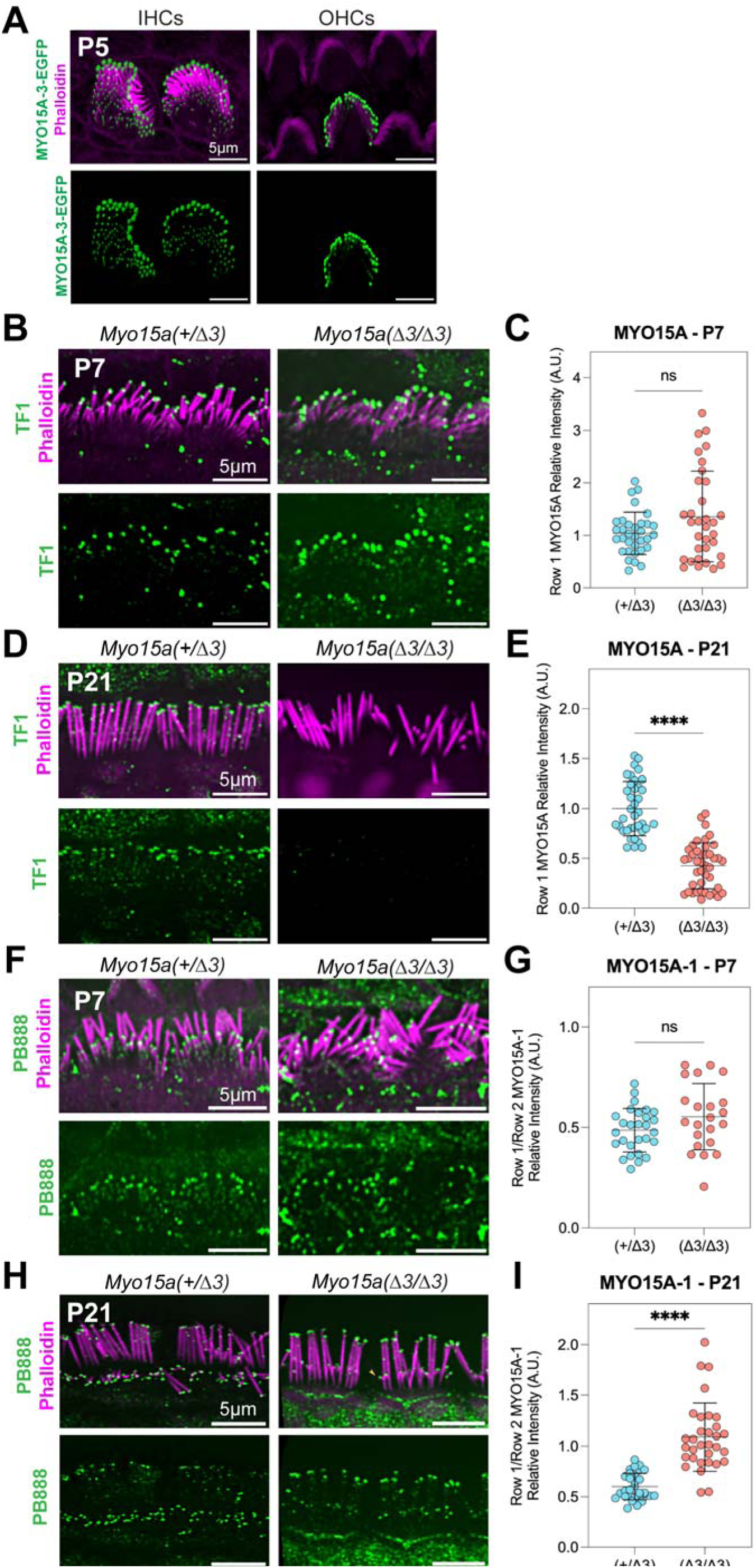
MYO15A-3 localizes to stereocilia hair cells. **(A)** Cochlear explants from postnatal day 5 (P5) of C57BL/6J mice were transfected to express the MYO15A-3 fused with EGFP (green). MYO15A-3 localizes at stereocilia tips of all rows in inner hair cells (IHCs) and outer hair cells (OHCs). **(B, D)** Stereocilia of *Myo15a^(+/^*^Δ*3)*^ and *Myo15a^(^*^Δ*3/*Δ*3)*^ mice at P7 and P21 mouse cochleae are labeled with MYO15A pan antibody (TF1) (green) and phalloidin (magenta). **(C, E)** Quantification of MYO15A localization at row 1 stereocilia at P7 and P21, determined by measuring the absolute fluorescence intensity at stereocilia tips. Each dot represents the fluorescence intensity of a single hair cell. This was measured by averaging the absolute fluorescence intensity at the tips of individual row 1 stereocilia. Data are mean ± SD of n=3 mice per genotype for P7 (3 female and 3 male) and P21 (all female) (total ∼37-42 hair cells per condition/genotype) with statistical significance of ***P<0.001 via Mann-Whitney U-test. **(F, H)** Immunofluorescence staining of MYO15A-1 using PB888 antibody in *Myo15a^(+/^*^Δ*3)*^ and *Myo15a^(^*^Δ*3/*Δ*3)*^ *a*t P7 and P21. **(G, I)** Quantification of MYO15A-1 localization was determined by measuring the relative fluorescent intensity ratio of Row1 to Row2 in *Myo15a^(+/^*^Δ*3)*^ and *Myo15a^(^*^Δ*3/*Δ*3)*^ at P7 and P21. Data are mean ± SD of n = 3 mice per genotype for P7 and P21 (4 female and 2 male) (total ∼22-32 hair cells per condition/genotype) with statistical significance of ****P<0.0001 via unpaired t-test. Scale bars: 5 μm.

The distribution of MYO15A-3 is similar to that reported for MYO15A-2 (Belyantseva et al. 2005; Fang et al. 2015), suggesting there may be a more complex cooperativity between these two isoforms in row 1 stereocilia. Whilst specific antibodies have detected endogenous MYO15A-1 (Fang et al. 2015) and MYO15A-3 (this study) in stereocilia, the localization of endogenous MYO15A-2 remains uncertain as there is no unique epitope that can be used to generate a truly specific antibody (Fig. 1B). One approach to localize endogenous MYO15A-2 has been to use pan-isoform MYO15A antibodies in MYO15A-1 knockout hair cells (Fang et al. 2015). This logic was premised upon wild-type hair cells expressing MYO15A-1 and MYO15A-2 only; thus, the pan-isoform antibody should specifically label residual MYO15A-2 in the knockout background (Fang et al. 2015). Our discovery that wild-type hair cells express three MYO15A isoforms falsifies this premise, as a pan-isoform antibody in MYO15A-1 knockout hair cells could potentially recognize either MYO15A-2 or MYO15A-3. The localization of MYO15A-2 therefore remains to be rigorously determined.

To help resolve this question, cochleae from *Myo15a^(^*^Δ*3/*Δ*3)*^ mice and *Myo15a^(+/^*^Δ*3)*^ littermates were labeled for IF using the previously characterized pan-isoform MYO15A antibody TF1 (Liang et al. 1999). We first examined TF1 immunoreactivity in hair cells at P7, when the expression is decreasing (Fig. 1G, K, N) and *Myo15a-3* expression is increasing (Fig. 1G, L, O). TF1 immunolabeling representing the summation of MYO15A-2 and MYO15A-3 isoforms was strongly detected at the tips of IHC row 1 stereocilia, and, to a lesser extent, at the tips of row 2 in control *Myo15a^(+/^*^Δ*3)*^ IHCs (Fig. 4B) as previously reported (Rzadzinska et al. 2004; Belyantseva, Boger, and Friedman 2003). TF1 labeling intensity at row 1 stereocilia tips was increased in *Myo15a^(^*^Δ*3/*Δ*3)*^ IHCs at P7 relative to controls (Fig. 4B, C). Although this trend was not statistically significant (Fig. 4C), it indicated that MYO15A-3 was not the predominant isoform at row 1 stereocilia tips at P7; we infer by exclusion that MYO15A-2 must therefore be present. We repeated this experiment at P21, when *Myo15a-3* expression is abundant in wild-type hair cells and *Myo15a-2* is downregulated (Fig. 1H, J, M). TF1 immunofluorescence was strongly localized to row 1 stereocilia tips in *Myo15a^(+/^*^Δ*3)*^ control IHCs at P21 and was significantly reduced in *Myo15a^(^*^Δ*3/*Δ*3)*^ knockout IHCs (Fig. 4D, E). We conclude that MYO15A-3 is the predominant species at row 1 stereocilia tips by P21, and that only residual quantities of MYO15A-2 remain. In addition, there is a switchover in myosin composition at row 1 stereocilia within the P7 – P21 window, with MYO15A-2 being largely replaced by MYO15A-3. This critical period (Stage IV) coincides with the final stage of stereocilia elongation, as row 1 the stereocilia reach their mature height (Krey et al. 2023, 2020; L G Tilney, Tilney, and DeRosier 1992; Lewis G. Tilney et al. 1986).

The increase in TF1 labeling observed at P7 in *Myo15a^(^*^Δ*3/*Δ*3)*^ IHCs was intriguing (Fig. 4C). Since TF1 has the potential to bind to all MYO15A isoforms (Fig. 1B), albeit with differing affinities, we hypothesized that the increase could be a result of another isoform being mis-localized to row 1 stereocilia. MYO15A-1 protein is detected in stereocilia from P4.5 onwards in mouse and primarily localizes to the shorter stereocilia (rows 2 and 3), which harbor active mechanoelectrical transduction (MET) channels (Fang et al. 2015; Krey et al. 2020; Beurg et al. 2009). To test if the loss of MYO15A-3 could affect the localization of MYO15A-1, we labeled control *Myo15a^(+/^*^Δ*3)*^ and *Myo15a^(^*^Δ*3/*Δ*3)*^ hair cells using the MYO15A-1 specific antibody PB888 (Fang et al. 2015). At P7, PB888 preferentially labeled the tips of row 2 and 3 stereocilia in both *Myo15a^(+/^*^Δ*3)*^ and *Myo15a^(^*^Δ*3/*Δ*3)*^ IHCs (Fig. 4F, G), indicating that MYO15A-1 localization was not perturbed by the loss of MYO15A-3. In P21 hair cells however, PB888 labeling significantly increased in the tallest stereocilia row 1 in *Myo15a^(^*^Δ*3/*Δ*3)*^ IHCs compared with control *Myo15a^(+/^*^Δ*3)*^ IHCs (Fig. 4H, I). Interestingly, despite MYO15A-1 being concentrated at row 1 stereocilia in *Myo15a^(^*^Δ*3/*Δ*3)*^ IHCs, this did not equate to labeling with TF1 (Fig. 4D), suggesting that the antibody either has a low affinity for MYO15A-1, or that the epitope was masked. In summary, our data reveals that the isoform composition of MYO15A changes at row 1 stereocilia tips through development; MYO15A-2 is present as stereocilia elongate, before transitioning to MYO15A-3 as they reach their terminal heights. The loss of MYO15A-3 in *Myo15a^(^*^Δ*3/*Δ*3)*^ IHCs also results in the mistargeting of MYO15A-1 towards row 1.

### MYO15A-3 can traffic components of the elongation complex along actin filaments

A central function of MYO15A-2 is to act as an intracellular trafficking motor that transports protein cargos along actin filaments to the stereocilia tip compartment (Belyantseva et al. 2005; Belyantseva, Boger, and Friedman 2003). Given that MYO15A-3 only differs from MYO15A-2 in their short N-terminal domain (Fig. 1B), we hypothesized that MYO15A-3 might similarly traffic along actin filaments and deliver EC proteins. To test this hypothesis, we used the ability of MYO15A-2’s ability to induce actin-based filopodial protrusions and traffic along them when over-expressed in immortalized cell lines (Belyantseva, Boger, and Friedman 2003; Belyantseva et al. 2005). Filopodial targeting of WHRN, EPS8, GNAI3, and GPSM2 all depend upon MYO15A-2 (Belyantseva et al. 2005; Delprat et al. 2005; Manor et al. 2011; Mauriac et al. 2017; Tadenev et al. 2019), and thus their filopodial tip localization can be used as a measure of trafficking efficiency.

As a positive control for our experiment, we transfected HeLa cells with plasmid DNA to over-express mCherry-tagged EPS8 and EGFP-tagged MYO15A-2 (Fig. 5A). MYO15A-2 and EPS8 fluorescence was co-localized at filopodia tips when visualized using confocal fluorescence microscopy (Fig. 5A), in agreement with previous work (Manor et al. 2011). Analysis of line scan fluorescence intensities along individual filopodia further showed that both MYO15A-2 and EPS8 signals were spatially correlated, indicating the presence of an interaction (Fig. 5D). As a negative control, we transfected HeLa cells with mCherry-EPS8 and EGFP-MYO10-HMM, a truncated MyTH4-FERM myosin that traffics on filopodia, but does not bind the EC proteins (Fig. 5B). In these cells, EPS8 did not accumulate at filopodia tips and line scans revealed that EPS8 and MYO10 signals were uncorrelated, indicating a lack of interaction (Fig. 5E). Finally, HeLa cells were transfected with mCherry-EPS8 and EGFP-MYO15A-3. In these cells, MYO15A-3 still co-accumulated at filopodia tips with EPS8 (Fig. 5C) and was also correlated in line scan analysis (Fig. 5F), indicating an interaction. We did note that MYO15A-3 targeted less efficiently to filopodia tips when compared with MYO15A-2 (Fig. 5A, C). To quantify these data, we used an algorithm developed previously (Bird et al. 2017) to measure the spatial correlation of two fluorophores along the filopodium (see Materials and Methods). Using this approach, we observed statistically significant spatial correlation of mCherry-EPS8 in 69.5 ± 8.4% (mean ± SD) of filopodia from cells expressing EGFP-MYO15A-3, versus 66.2 ± 9.0% of filopodia from cells expressing EGFP-MYO15A-2, and 9.5 ± 1.6% of filopodia from cells expressing EGFP-MYO10-HMM (Fig. 5G). We conclude that MYO15A-3 is able to bind and traffic EPS8 along filopodia actin filaments, similar to MYO15A-2.

**Figure 5.**
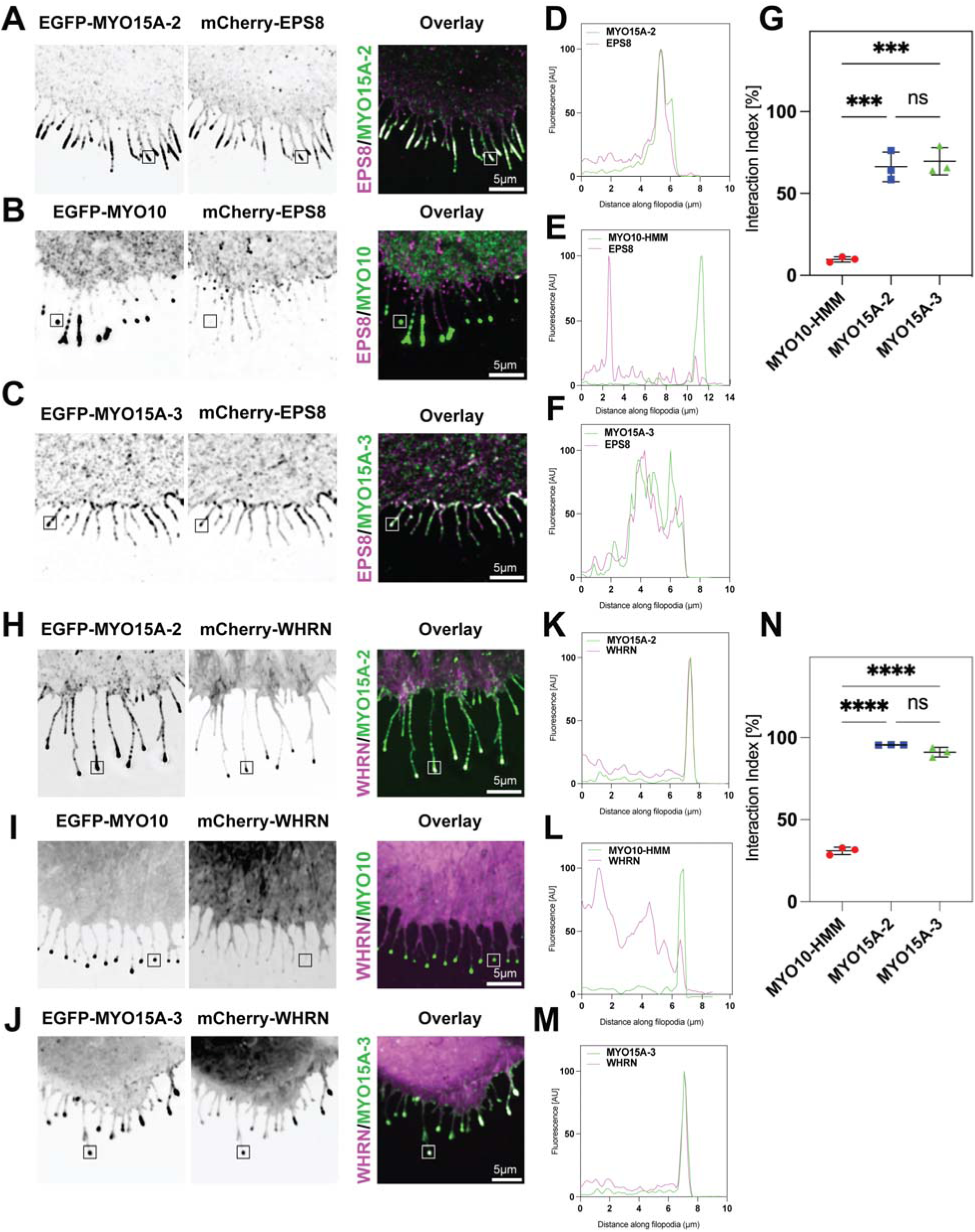
MYO15A-3 traffics the elongation complex protein in filopodia. HeLa cells were transfected with plasmids encoding for protein fusion proteins, either EGFP-MYO15A-2 (positive control), EGFP-MYO15A-3 or EGFP-MYO10 (negative control), and co-transfected with either **(A-C)** mCherry-EPS8 or (H-J) mCherry-WHRN to detect endogenous protein. Cells were imaged using a spinning disk confocal microscope to investigate the localization and interaction of these proteins. The intensity of EGFP-Myosin and mCherry-EPS8/WHRN was measured blinded at multiple filopodia tips (∼300 filopodia from n=3 independent transfections). **(D-F, K-M)** The molecular interaction between the myosin and the EC proteins was determined using Pearson’s r coefficient, calculated along the filopodia shaft to measure the fluorescence correlation between the bait and prey. **(G, N)** Each data point represents the average interaction index from a single experiment (3 independent determinations) with a total of 30 cells per condition. The data are presented as mean ± SD with statistical significance determined using One-way ANOVA. *** P<0.001, **** P<0.0001. Scale bars: 5 µm.

The potential interaction of WHRN with MYO15A-3 was tested using a similar methodology. In HeLa cells over-expressing mCherry-WHRN and EGFP-MYO15A-2 as a positive control, there was robust co-accumulation of WHRN and MYO15A-2 at filopodia tips (Fig. 5H, K), similar to previous work (Belyantseva et al. 2005). In cells expressing mCherry-WHRN and EGFP-MYO10-HMM as a negative control, WHRN did not accumulate at filopodia tips despite MYO10 trafficking to this location, indicating the absence of interaction (Fig. 5I, L). In cells expressing mCherry-WHRN and EGFP-MYO15A-3, we again observed co-localization of MYO15A-3 and WHRN at filopodia tips (Fig. 5J, M). Quantification of line scans taken along individual filopodia confirmed the statistically significant spatial correlation of mCherry-WHRN in 91 ± 3.0% (mean ± SD) of filopodia in cells expressing EGFP-MYO15A-3, 95.6 ± 0.1% of filopodia in cells expressing EGFP-MYO15A-2, and 30.8 ± 2.3% of filopodia in cells expressing EGFP-MYO10-HMM (Fig. 5N). We conclude that MYO15A-3 can bind and traffic WHRN along filopodia actin filaments. In summary, our *in vitro* experiments show that MYO15A-3 is similar to MYO15A-2 and can bind and traffic at least two core components of the elongation complex, WHRN and EPS8.

### MYO15A-3 traffics the elongation complex along stereocilia in postnatal hair cells

Elongation complex (EC) proteins are necessary for establishing stereocilia row identity and elongation during hair bundle development (Belyantseva et al. 2005; Manor et al. 2011; Mauriac et al. 2017; Tadenev et al. 2019); these proteins persist at row 1 stereocilia tips in adult hair cells (Hartig et al. 2024; Zampini et al. 2011; Belyantseva et al. 2005). Given MYO15A-3 localization to row 1 stereocilia in adult hair cells, and its trafficking of EPS8 and WHRN for *in vitro* heterologous cells, we hypothesize that MYO15A-3 delivers EC proteins in postnatal hair cells after MYO15A-2 is transcriptionally downregulated. To test this hypothesis, we labeled for EPS8 and WHRN immunofluorescence in *Myo15a^(+/^*^Δ*3)*^ and *Myo15a^(^*^Δ*3/*Δ*3)*^ IHCs at P4 and P21, ages when the composition of row 1 stereocilia tips was primarily MYO15A-2 or MYO15A-3, respectively.

At P4, EPS8 is primarily concentrated at row 1 stereocilia tips in both control *Myo15a^(+/^*^Δ*3)*^ and mutant *Myo15a^(^*^Δ*3/*Δ*3)*^ IHCs (Fig. 6A), and there is no statistically significant difference in row 1 fluorescence intensities between genotype (Fig. 6B). Similarly, WHRN immunofluorescence is concentrated at row 1 stereocilia tips in both *Myo15a^(+/^*^Δ*3)*^ and *Myo15a^(^*^Δ*3/*Δ*3)*^ IHCs (Fig. 6C), and quantification revealed no significant difference between fluorescence intensities at row 1 tips (Fig. 6D). WHRN immunofluorescence is also concentrated at the base of stereocilia in both genotypes (Fig. 6C), where it forms part of the ankle-link complex that is not dependent upon MYO15A (Michalski et al. 2007). We conclude that MYO15A-3 is not required for the initial targeting of EPS8 and WHRN to row 1 stereocilia tips at P4, and that MYO15A-2 is sufficient to deliver these proteins. We next examined EPS8 and WHRN in both *Myo15a^(+/^*^Δ*3)*^ and *Myo15a^(^*^Δ*3/*Δ*3)*^ IHCs at P21. In control *Myo15a^(+/^*^Δ*3)*^ IHCs, both EPS8 (Fig. 6E) and WHRN (Fig. 6G) are concentrated at the tips of row 1 stereocilia (Fig. 6E, G). In mutant *Myo15a^(^*^Δ*3/*Δ*3)*^ IHCs however, the fluorescence of EPS8 (Fig. 6E) and WHRN (Fig. 6G) were significantly reduced compared with control *Myo15a^(+/^*^Δ*3)*^ IHCs (Fig. 6F, H). We conclude that the correct targeting of EPS8 and WHRN to row 1 stereocilia tips does not initially require MYO15A-3 at P4, consistent with MYO15A-2 being the major species at this age. By P21 however, EPS8 and WHRN trafficking must become dependent upon MYO15A-3. These findings suggest that during postnatal hair bundle maturation, the transport of EC protein transitions from utilizing MYO15A-2 to MYO15A-3.

**Figure 6.**
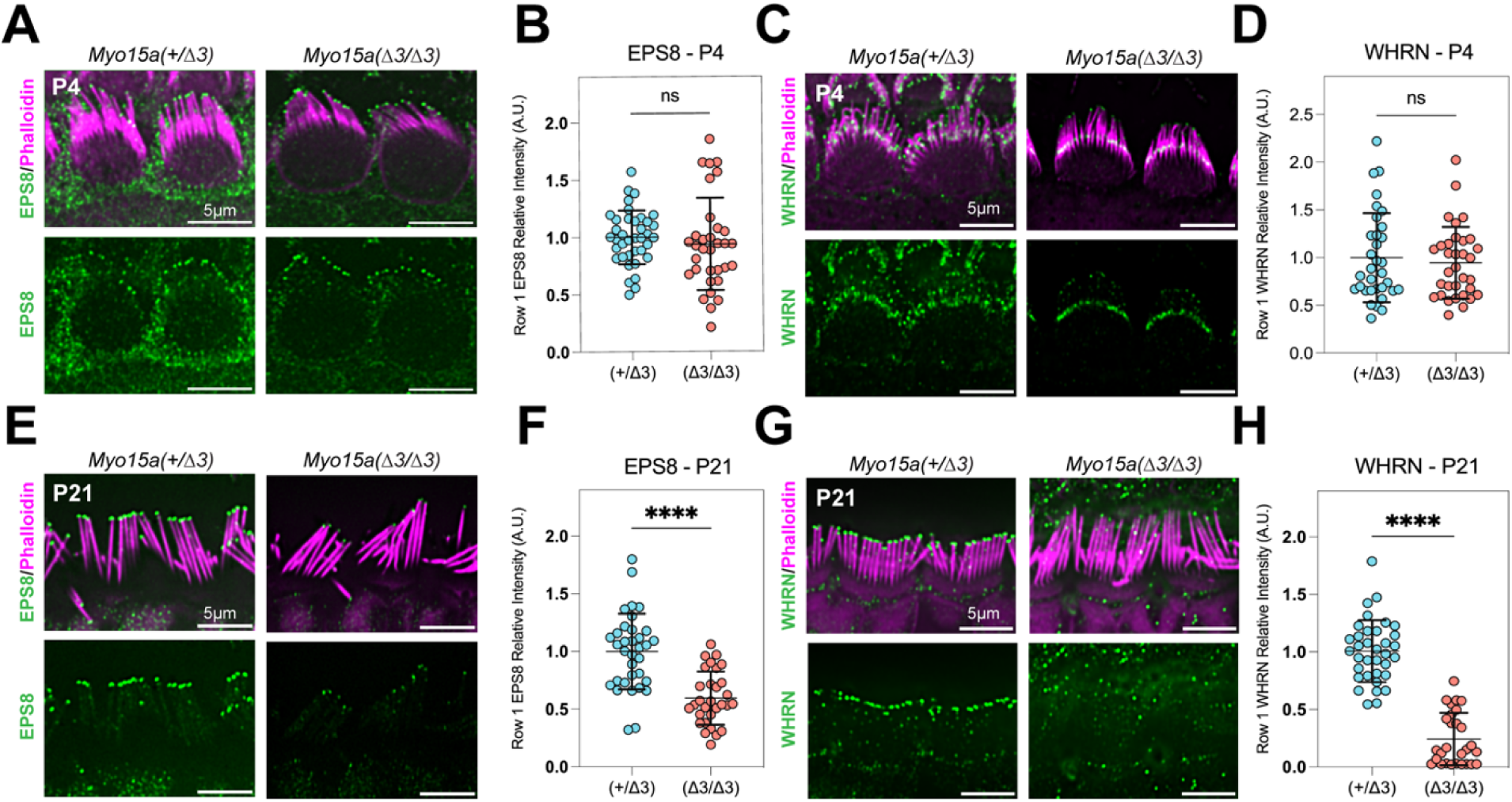
MYO15A-3 is not required for the elongation complex trafficking in hair cells at P4, but traffics the elongation complex at P21. **(A, C)** Confocal images of hair cells from cochleae of *Myo15a^(+/^*^Δ*3)*^ and *Myo15a^(^*^Δ*3/*Δ*3)*^ mice at postnatal day 4 (P4) immunolabeled for the EPS8/ WHRN (green). The hair cells were counterstained with phalloidin (magenta) to visualize the actin-rich stereocilia bundles. **(B, D)** Quantification of EPS8 and WHRN relative intensity at the stereocilia tips. Each dot represents the fluorescence intensity of a single hair cell. This was measured by averaging the absolute fluorescence intensity at the tips of individual row 1 stereocilia. Data are mean ± SD for n=3 mice per genotype (total ∼33-38 hair cells per condition/genotype). The absolute fluorescence for EPS8/WHRN was not significantly different between genotypes, as determined via the Mann-Whitney U-test to measure EPS8 level and an unpaired t-test for WHRN. **(E, G)** Confocal images of hair cells from cochleae of *Myo15a^(+/^*^Δ*3)*^ and *Myo15a^(^*^Δ*3/*Δ*3)*^ mice at postnatal day 21 (P21) immunolabeled for the EPS8/ WHRN (green). **(F, H)** Quantification of EPS8 and WHRN relative intensity at the stereocilia tips. Data are mean ± SD for n=3 mice per genotype (total ∼28-35 hair cells per condition/genotype) with statistical significance of **** P<0.0001 via Mann-Whitney U-test to measure WHRN and an unpaired t-test for EPS8. Scale bars: 5 µm.

### Postnatal maintenance of hair cell stereocilia architecture requires MYO15A-3

The disruption of EC trafficking in *Myo15a^(^*^Δ*3/*Δ*3)*^ mice was evident by P21, and this corresponded with hearing loss that rapidly progressed in severity from P17 to P30. To explore the ultrastructural basis for hearing loss, we used scanning electron microscopy (SEM) to study hair bundle morphology of mutant *Myo15a^(^*^Δ*3/*Δ*3)*^ and normal hearing *Myo15a^(+/^*^Δ*3)*^ littermates. We imaged *Myo15a^(^*^Δ*3/*Δ*3)*^ hair cells at P7 and P21 to reflect key timepoints when the EC protein complex was either present (P7), or absent (P21), respectively. At P7, IHC hair bundles from *Myo15a^(^*^Δ*3/*Δ*3)*^ mice had a normal staircase architecture and were qualitatively indistinguishable from *Myo15a^(+/^*^Δ*3)*^ littermate controls (Fig. 7A, G). There were no statistically significant differences between genotypes in IHCs row 1 stereocilia widths (Fig. 7C) at different positions relative to the tip. Row 2 stereocilia widths were also equivalent between genotypes in IHCs (Fig. 7D). This data indicates MYO15A-3 is not required for the initial assembly of the hair bundle, consistent with the normal localization of the EC in *Myo15a^(^*^Δ*3/*Δ*3)*^ mice at P4.

**Figure 7.**
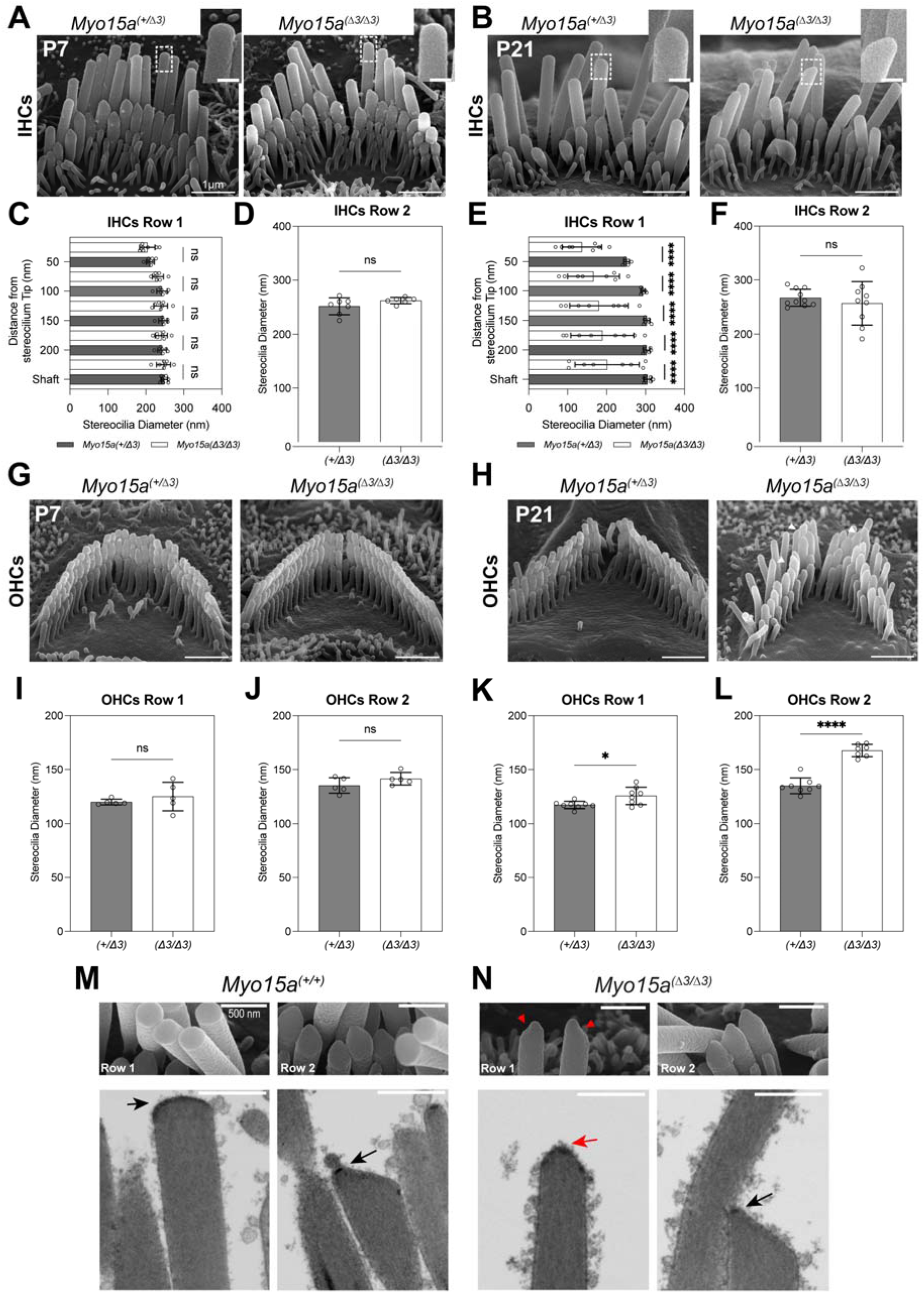
Stereocilia degenerate in *Myo15a^(^*^Δ*3/*Δ*3)*^ IHCs and OHCs postnatally. Scanning electron microscopy (SEM) images of *Myo15a^(+/^*^Δ*3)*^ and *Myo15a^(^*^Δ*3/*Δ*3)*^ IHCs and OHCs stereocilia bundles at **(A, G)** P7 and **(B, H)** P21. **(C, E)** Stereocilia diameters from the first row of IHCs measured at the shaft and multiple positions near the tip in P7 *Myo15a^(+/^*^Δ*3)*^ (Dark gray n=7 hair cells) and *Myo15a^(^*^Δ*3/*Δ*3)*^ mice (white n=7 hair cells). At P21 from *Myo15a^(+/^*^Δ*3)*^ (dark gray, n=7 hair cells) and *Myo15a^(^*^Δ*3/*Δ*3)*^ (white, n=9 hair cells) mice. **(D, F)** The diameter of row 2 IHCs at P7 (n=6-7 hair cells) and P21 (n=∼10 hair cells) measured at 200 nm from the tip. **(I, J)** The diameter of row 1 (n=5 hair cells) and row 2 OHCs (n=8-9 hair cells) at P7 measured 200 nm from the tip. **(K, L)** The same measurement was done to determine the diameter of row 1 (n=8 hair cells) and row 2 OHCs (n=8 hair cells) at P21 for both genotypes. Data are mean ± SD. * P<0.05, **** P<0.0001, with statistical analysis for row 1 IHC diameter conducted using One-way ANOVA, Šidák, and Dunn’s multiple comparison tests, and unpaired t-tests for comparing the diameters of row 1 OHCs and row 2 IHCs. Scale bar: 1 µm. Adult (P21) stereocilia from row 1 and row 2 were imaged using focused ion beam scanning electron microscopy (FIB-SEM) with 20 nm serial sectioning to view tip densities. (M) *Myo15a^(+/+)^* and (N) *MYO15a^(^*^Δ*3/*Δ*3)*^. Black and red arrows indicate normal and abnormal tip densities, respectively. Bottom panels are minimum intensity Z-projections of five consecutive slices, or 100nm thickness. Scale bars: 500nm, 1µm.

In contrast, a range of degenerative changes to the hair bundle and individual stereocilia were observed in *Myo15a^(^*^Δ*3/*Δ*3)*^ hair cells at P21. In IHCs, there was a generalized thinning of row 1 stereocilia (Fig. 7B), which was pronounced within 50 nm of the stereocilia tip (Fig. 7E). Row 1 stereocilia tips were misshapen in *Myo15a^(^*^Δ*3/*Δ*3)*^ IHCs (Fig. 7B, insets) and lacked the hemi-spherical cap that was evident in *Myo15a^(+/^*^Δ*3)*^ controls (Fig. 7B, insets). Analysis of row 1 stereocilia widths revealed a statistically significant reduction in diameter 50 nm from the tip, from 251 ± 7 nm in *Myo15a^(+/^*^Δ*3)*^ to 137 ± 50 nm in *Myo15a^(^*^Δ*3/*Δ*3)*^ IHCs (Fig. 7E). No statistically significant changes were observed in row 2 stereocilia widths in either *Myo15a^(+/^*^Δ*3)*^ or *Myo15a^(^*^Δ*3/*Δ*3)*^ IHCs (Fig. 7F). In conclusion, MYO15A-3 is not required for hair cells to assemble stereocilia into a morphologically normal hair bundle but instead is necessary to maintain the size and stability of the tallest row 1 stereocilia.

### The delivery of elongation complex proteins by MYO15A-3 is necessary for maintenance of the stereocilia tip-density

A prominent feature of *Myo15a^(^*^Δ*3/*Δ*3)*^ hair cells was the structural degeneration at the tips of row 1 stereocilia, which correlated with the loss of MYO15A-3, EPS8, and WHRN from the same location. We hypothesized that MYO15A-3 is required to maintain the tip density; which is a protein-dense osmiophilic structure at row 1 stereocilia tips that is thought to contain the regulatory machinery controlling actin polymerization (Furness and Hackney 1985; Rzadzinska et al. 2004). EC proteins and MYO15A are proposed to constitute the tip density by forming a biomolecular condensate via liquid-liquid phase separation (LLPS) (Lin et al. 2021; Shi et al. 2022). Tip densities are absent in mutant *Myo15a shaker-2* hair cells (Rzadzinska et al. 2004), and significantly disrupted in *Whrn* knockout hair cells (Mogensen, Rzadzinska, and Steel 2007), showing that both MYO15A and EC proteins contribute to formation of the tip-density during stereocilia development. Using focused ion beam scanning electron microscopy (FIB-SEM), we analyzed the structure of IHC row 1 and row 2 stereocilia tips from *Myo15a^(^*^Δ*3/*Δ*3)*^ and littermate *Myo15a^(+/+)^* controls at P21. Row 1 stereocilia in *Myo15a^(+/+)^* IHCs had a well-defined electron-dense band along the edge of their characteristic oblate tip (Fig. 7M). FIB-SEM also revealed the lower tip-link density (LTLD) within the prolate shaped tips of row 2 stereocilia (Fig. 7M). In contrast, the row 1 stereocilia tip density in *Myo15a^(^*^Δ*3/*Δ*3)*^ IHCs was discontinuous and patchy (Fig. 7N). This corresponded with stereocilia thinning observed on SEM. The LTLD at the tips of row 2 stereocilia was normal in *Myo15a^(^*^Δ*3/*Δ*3)*^ IHCs, consistent with the architecture of row 2 stereocilia being unaffected at this age (Fig. 7F). In conclusion, our data supports a model where MYO15A-3 traffics and delivers the EC to row 1 stereocilia tips, and that this complex is necessary for the continued postnatal integrity of stereocilia tip densities and hair bundle architecture.

## DISCUSSION

Our study reveals unexpected complexity in the mechanisms that MYO15A uses to shape hair cell stereocilia and uncovers an additional etiology for human hearing loss DFNB3. We found that a previously uncharacterized isoform, MYO15A-3, maintains the postnatal structure of mechanosensory cochlear hair cell stereocilia and is necessary for hearing. Mutant mice with a specific deletion of *Myo15a-3* exhibited rapidly progressing hearing loss that caused complete deafness by P30. Hair bundles in *Myo15a^(^*^Δ*3/*Δ*3)*^ mutants were indistinguishable from wild-type controls at P7 but developed a unique stereocilia pathology by P21, mirroring the postnatal degeneration of auditory function. Stereocilia degeneration was coupled to the loss of key elongation complex proteins EPS8 and WHRN, which that depended upon MYO15A-3 for stereocilia trafficking and are necessary for stereocilia assembly. We conclude that a complex of MYO15A-3, EPS8, and WHRN is required for postnatal stereocilia maintenance and hearing. Our data implies that in addition to their well characterized role in stereocilia development, elongation complex proteins are also necessary to maintain stereocilia architecture into adult life. Interestingly, the trafficking of elongation complex proteins involved a handoff between MYO15A isoforms; EPS8 and WHRN were initially delivered by MYO15A-2 during stereocilia elongation, then transitioned to MYO15A-3-based trafficking as stereocilia attained their mature size. Thus, normal hearing function depends upon an overlapping network of three MYO15A isoforms that cooperate to establish and fine-tune stereocilia architecture throughout life. The interleaved, non-redundant activities of MYO15A isoforms represent three distinct molecular pathologies that underlie human hearing loss DFNB3.

The initial discovery that MYO15A is required for stereocilia growth was based on studies using the shaker 2 (*Myo15a^sh2^*) mouse harboring an inactivating mutation in the MYO15A ATPase domain (Probst et al. 1998); however, the *shaker 2* mutation is common to all three isoforms expressed by hair cells (Fig. 1B) and cannot discriminate between the function of MYO15A-1 versus MYO15A-2, or MYO15A-3. More recently, mouse mutants with isoform-selective deletions have teased out additional functions for MYO15A. Selective deletion of MYO15A-1 (*Myo15a*^Δ*N*^) allows stereocilia elongation but causes degeneration of mechano-transducing row 2 and 3 stereocilia from ∼P7 onwards (Fang et al. 2015; Nal et al. 2007). These data also imply that MYO15A-2 and or MYO15A-3 alone must be necessary for stereocilia elongation. Here, we show that *Myo15a(*^Δ*3/*Δ*3*^) hair bundles develop normally up until at least P7, demonstrating that MYO15A-3 is not required for initial stereocilia development. We conclude that MYO15A-2 is likely sufficient for stereocilia development *in vivo*, expanding upon previous *in vitro* findings that exogenous EGFP-tagged MYO15A-2 can rescue hair cell stereocilia elongation in *shaker 2* explant cultures (Belyantseva et al. 2005). We cannot exclude that the activities of MYO15A-2 and MYO15A-3 are redundant during stereocilia elongation. However, our analysis of *Myo15a* transcripts using qRT-PCR and BaseScope are not consistent with this interpretation; *Myo15a-3* expression increased postnatally, whereas *Myo15a-2* declined neonatally as stereocilia reach their adult height.

Several hundred pathological variants linked to DFNB3 have been identified across the *MYO15A* locus (Rehman et al. 2016; J. Zhang et al. 2019; Miyoshi et al. 2024); this includes variants in exon 2 specific to MYO15A-1 and in the ATPase and tail domain-encoding exons that are common to all three isoforms. Pathogenic variants are yet to be reported in exon 1 of MYO15A-3, however a deeply intronic non-coding variant at *MYO15A* c.3609(+985)G>A was identified in multiple pedigrees segregating DFNB3 deafness that was computationally predicted to eliminate splice donor activity of MYO15A-3 exon 1 (Abu Rayyan et al. 2020). These data suggest that our approach of deleting exon 1 using CRISPR/Cas9 in mouse reflects a clinically relevant cause of DFNB3 deafness. The identification of variants within MYO15A-3 exon 1 should now be included in genetic screening for patients with suspected hereditary hearing loss. Our study provides an updated framework to assess the pathogenicity of novel variants potentially targeting MYO15A-3 and further expands the phenotypic spectrum of DFNB3.

The structural phenotype of MYO15A-3 null hair cells is unique amongst *Myo15a* mutant mouse models, with disruption of the row 1 tip density that correlated with row 1 stereocilia thinning. OHCs had a similar structural disruption to row 1 stereocilia tips, but we did not detect row 1 stereocilia thinning at the timepoints analyzed. The *Myo15a^(^*^Δ*3/*Δ*3)*^ hair cell phenotype is unlike the *Myo15a shaker 2*, *shaker 2J* and *jordan* mouse models that have short stereocilia and additional stereocilia rows (Probst et al. 1998; Anderson et al. 2000; Moreland et al. 2025). *Myo15a^(^*^Δ*3/*Δ*3)*^ hair cells instead assembled a structurally normal hair bundle, up until at least P7, that was similar to *Myo15a^(^*^Δ*N/*Δ*N)*^ hair cells, in agreement with both MYO15A-3 and MYO15A-1 being dispensable for stereocilia growth (Fang et al. 2015). The *Myo15a^(^*^Δ*3/*Δ*3)*^ *and Myo15a^(^*^Δ*N/*Δ*N)*^ hair cell phenotypes diverge after P7; with structural degeneration observed in *Myo15a^(^*^Δ*3/*Δ*3)*^ row 1 stereocilia, versus mechanosensitive rows 2 and 3 in *Myo15a^(^*^Δ*N/*Δ*N)*^ hair cells. This difference in postnatal degeneration is mirrored by the distribution of MYO15A proteins, with MYO15A-3 concentrated at row 1 stereocilia tips and MYO15A-1 at row 2 and 3 tips. Our results delineate three essential phases of MYO15A activity; MYO15A-2 is required for initial stereocilia growth, whilst MYO15A-1 and MYO15A-3 contribute to distinct postnatal mechanisms that maintain stereocilia architecture.

How does MYO15A-3 control the architecture of row 1 stereocilia? We found that MYO15A-3 could traffic EPS8 and WHRN along filopodia in HeLa cells *in vitro*, and that EPS8 and WHRN were significantly reduced at row 1 stereocilia tips in MYO15A-3 null hair cells *in vivo*. EPS8 and WHRN did not require MYO15A-3 to localize correctly at P4 but required MYO15A-3 by P21; this age-selective dependence mirrored the transcriptional switch from MYO15A-2 to MYO15A-3 in postnatal hair cells. We propose that EPS8 and WHRN are delivered to neonatal row 1 stereocilia by MYO15A-2, before a postnatal handover to MYO15A-3 based trafficking. This model explains why stereocilia architecture was initially preserved in *Myo15a^(^*^Δ*3/*Δ*3)*^ neonatal hair cells through MYO15A-2 driven trafficking of the EC; degeneration results from the failed handover to MYO15A-3, disrupting the postnatal delivery of EC proteins. Stereocilia degeneration in postnatal *Myo15a^(^*^Δ*3/*Δ*3)*^ hair cells resulted from the concomitant loss of MYO15A-3, EPS8 and WHRN. A limitation of our study is that we cannot dissect the individual contributions of these proteins; however, our results support the complex of MYO15A-3, EPS8 and WHRN maintaining row 1 stereocilia. EPS8 and WHRN are individually required for stereocilia elongation, and along with GPSM2 and GNAI3, establish the molecular identity of row 1 stereocilia during hair bundle morphogenesis (Manor et al. 2011; Mburu et al. 2003; Belyantseva et al. 2005; Mauriac et al. 2017; Tadenev et al. 2019). Curiously, EPS8, WHRN, GPSM2 and GNAI3 all remain strongly concentrated at row 1 stereocilia tips in adult hair cells (Zampini et al. 2011; Belyantseva et al. 2005; Hartig et al. 2024). Postnatal inactivation of *Gpsm2* results in shortening of row 1 stereocilia with degeneration focused within 0.5 µm of the stereocilia tip (Hartig et al. 2024). This phenocopies row 1 stereocilia in *Myo15a^(^*^Δ*3/*Δ*3)*^ hair cells and suggests that MYO15A-3 and GPSM2 function in a common molecular pathway to maintain adult stereocilia architecture. GPSM2, along with GNAI3, potentiates the liquid-liquid phase separation (LLPS) of WHRN, EPS8 and MYO15A to form biomolecular condensates that are proposed to form the tip density complex *in vivo* (Lin et al. 2021; Shi et al. 2022). The simultaneous loss of WHRN, EPS8 and MYO15A-3 can therefore be expected to abolish condensate formation; consistent with this, we observed disruption to the tip density in *Myo15a^(^*^Δ*3/*Δ*3)*^ hair cells. Actin polymerization occurs continuously at the tips of row 1 stereocilia (Narayanan et al. 2015; D.□S. Zhang et al. 2012) and our data suggests that MYO15A-3 and the EC are necessary to regulate this activity and tune row 1 stereocilia size.

In parallel with the loss of MYO15A-3, WHRN and EPS8 from row 1 stereocilia, we also detected the mistargeting of other MYO15A isoforms in *Myo15a^(^*^Δ*3/*Δ*3)*^ IHCs. In wild-type IHCs, MYO15A-1 is normally restricted to mechanosensitive shorter stereocilia rows 2 and 3 from ∼P4.5 onwards in close proximity to the MET apparatus (Miller, Wang, and Grillet 2024; Fang et al. 2015). MYO15A-1 was correctly targeted in *Myo15a^(^*^Δ*3/*Δ*3)*^ IHCs at P7 but redistributed to row 1 stereocilia tips by P21. Consistent with MYO15A-1 not delivering EPS8 and WHRN in wild-type hair cells (Fang et al. 2015), the mistargeting of MYO15A-1 could not restore EPS8 or WHRN in row 1. MYO15A-1 can redistribute to the lateral tips of row 1 stereocilia in *Tmie(-/-)* mutants lacking MET currents (Krey et al. 2020), and infrequently in wild-type hair cells, presumably due to sporadic hair bundle and tip-link trauma (Fang et al. 2015). Retargeting in *Myo15a^(^*^Δ*3/*Δ*3)*^ IHCs was distinct from these examples and MYO15A-1 formed a robust cap at row 1 stereocilia tips, similar to that revealed using pan-MYO15A antibodies (Rzadzinska et al. 2004). The mis-localization of MYO15A-1 was only observed in the absence of MYO15A-3 and following the transcriptional downregulation and loss of MYO15A-2 from hair cells after P7. We speculate that MYO15A-1 competes for the same binding sites as MYO15A-2 and MYO15A-3 within stereocilia but with a lower affinity that normally prevents its association. This affinity-based exclusion model is consistent with GPSM2 / GNAI3 refining the initial targeting of MYO15A-2 to row 1 stereocilia (Tadenev et al. 2019), leaving shorter mechanosensitive rows accessible to MYO15A-1. How the mistargeting of MYO15A-1 contributes to hearing loss and the degenerative row 1 phenotype in *Myo15a^(^*^Δ*3/*Δ*3)*^ hair cells is an important area of future investigation.

A fascinating question remains; why do hair cells use MYO15A-2 and MYO15A-3 as two separate isoforms to traffic EC proteins in stereocilia? Our results argue that MYO15A-2 drives initial stereocilia development, with its abundance waning by P7 as stereocilia enter their terminal elongation phase (Krey et al. 2023; Lewis G. Tilney et al. 1986; L G Tilney, Tilney, and DeRosier 1992). Thus, a gradual handover to MYO15A-3 occurs as stereocilia attain their mature architecture. Both isoforms are similar and only differ by the 6 kDa N-terminal domain unique to MYO15A-3 (Rehman et al. 2016). We hypothesize that the 6 kDa N-terminal domain must supply a unique and irreplaceable function. N-terminal domains are a general feature of the myosin superfamily and can modulate the ATPase force generating mechanochemical cycle. For example, the N-terminal kinase domain of MYO3A auto-phosphorylates and regulates the ATPase domain (Quintero et al. 2010; Dosé et al. 2008). Other myosin motors, including MYO1B, MYO1C and MYO7A have non-catalytic N-terminal domains that modulate the ATPase mechano-chemical cycle to alter force sensing and force generation (Holló et al. 2023; Greenberg et al. 2015; Li et al. 2020). In HEK293 cells, both MYO15A-2 and MYO15A-3 trafficked EPS8 and WHRN along filopodia, suggesting broadly equivalent activity; though, we cannot exclude modulation of the ATPase that could alter other properties, such as: actin affinity, velocity, duty ratio, and other enzymatic parameters.

The ATPase domain of MYO15A can catalyze the nucleation of actin filaments (Gong et al. 2022), a property required for stereocilia elongation (Moreland et al. 2025), along with modulating force generation. It is yet unknown how the 6 kDa N-terminal domain of MYO15A-3 alters nucleation activation of the ATPase; however, we speculate that if MYO15A-3 were unable to nucleate actin filaments, the shift from MYO15A-2 (nucleation competent) to MYO15A-3 (nucleation incompetent) could restrict stereocilia growth as row 1 attain their mature size. In this way, the MYO15A-2:MYO15A-3 ratio could tune activity of the myosin motor ensemble at row 1 stereocilia tips, similar to MYO7A at the UTLD where the ratio of splice isoforms is proposed to control tip link tension (Li et al. 2020). Though speculative, our model may explain why *Myo15a-2* is re-expressed in a subset of hair cells at P60. Supporting its role in early stereocilia elongation, *Myo15a-2* was expressed at P0, but downregulated and undetectable by P14 using BaseScope. By P60 however, *Myo15a-2* was reproducibly detected in ∼10% of IHCs and OHCs, suggesting that MYO15A-2 and MYO15A-3 can both occupy row 1 stereocilia in adult hair cells. Actin filaments continually turnover at the tips of row 1 stereocilia in adult hair cells *in vivo* (Narayanan et al. 2015), and this activity is influenced by GPSM2 (Hartig et al. 2024) and the MYO15A-3 complex with EPS8 and WHRN, as our present study indicates. Altering the relative ratios of MYO15A-2 and MYO15A-3 isoforms postnatally may alter force generation or actin polymerization at row 1 and fine-tune stereocilia size. In conclusion, our results reveal a MYO15A based mechanism that is necessary for the continued structural maintenance of row 1 stereocilia and highlights three separate activities of MYO15A that establish and orchestrate stereocilia architecture. With the advent of clinical genetic therapies to treat hereditary hearing loss in humans (Qi et al. 2024; Lv et al. 2024), our study establishes the essential spectrum of MYO15A isoform activities that must be corrected to successfully treat DFNB3.

## MATERIALS AND METHODS

### Generation of the *Myo15a-***Δ***3* allele using CRISPR-Cas9 mutagenesis

A mutant allele deleting exon 1 of *Myo15a-3* was created using CRISPR-mediated non-homologous end joining (NHEJ) on a C57BL/6N genetic background. Briefly, two guide RNAs (gRNA) flanking exon 1 (mm10 chr11:60,480,418–60,480,621) were chosen using CRISPRscan (www.CRISPRscan.org) and synthesized with T7 *in vitro* transcription as described (Varshney et al. 2015). Guide RNA cleavage activity was assessed *in vitro* by incubating a PCR product encompassing the target sites with SpCas9 (PNA Bio, Thousand Oaks, CA) and analyzing products on a 2% agarose gel. Next, gRNA functionality was validated via Surveyor assay in an immortalized mouse embryonic fibroblast (MEF) cell line with a tetracycline-inducible Cas9 expression cassette. Following confirmation of efficient target cleavage, the two gRNAs (Myo15a_gR1: 5’-GGG TGG CCC TTG GAG GAG G-3’ and Myo15a_gR5: 5’-GGC AAG CAC GTG AGC CGA T-3’) were mixed with SpCas9 and microinjected into C57BL/6N mouse zygotes as described (H. Wang et al. 2013). Genomic DNA from six pups was screened by PCR (primers: 5’-GGA CTA GAC ATG AGA GTA GAG GCA A-3’ and 5’-CTG TGA GCA CAC ACG TGT GCA GCC CT-3’). Two founders generated a 326 bp amplicon that was Sanger sequenced to show a 331 bp deletion compared to the wild-type (657 bp) allele. This deletion removed exon 1 entirely. Germline transmission of the *Myo15a-*Δ*3* allele was confirmed in F1 progeny, and mice were subsequently backcrossed to C57BL/6J to establish the *Myo15a-*Δ*3* mutant strain. Mouse mutagenesis was approved by the NIH IACUCs (#NEI-626 & #NINDS-1263-20). All animal experiments were approved by the University of Florida Institutional Animal Care and Use Committee (protocol # IACUC202200000513).

### Genotyping of the *Myo15a-*Δ*3* allele

Routine genotyping was performed using a multiplex hydrolysis assay on a QuantStudio 6 Flex Real-Time PCR System (Applied Biosystems). The genotyping strategy was designed to differentiate between the wild-type (WT) exon 1 of *Myo15a-3* and the deletion mutant allele present in *Myo15a-*Δ*3* mutant mice in a single reaction. To amplify the wild-type allele, a 6-FAM 5’-labeled probe and flanking primers, were designed to anneal within the sequence deleted in the *Myo15a(*Δ*3)* allele. To detect the mutant allele, a HEX 5’-labeled probe was positioned to span the CRISPR-generated recombination scar, with primers located in adjacent wild-type sequence flanking exon 1. All probes and primers (Table 1) were synthesized by Integrated DNA Technologies (Coralville, IA). Genomic tail DNA was combined with all primers, FAM and HEX-labeled probes, and a qPCR master mix (KAPA Probe Fast, #KR0397, Roche) following the manufacturer’s protocol. The thermal profile for amplification was 95 °C (3 m, followed by 40 cycles of 95 °C (3 s and 60 °C (30 secs). For each reaction, an XY scatter plot was generated, with FAM fluorescence (wild-type) on the abscissa, and HEX fluorescence (mutant) on the ordinate. Each data point represented the cycle-by-cycle amplification of matched FAM / HEX pair from a single sample. The genotype was interpreted from a line of best fit, with a horizontal line indicating homozygous wild-type, a vertical line indicating homozygous mutant, and diagonal indicating a heterozygous genotype. A supplementary python script was written to automate this analysis.

**Table 1.**
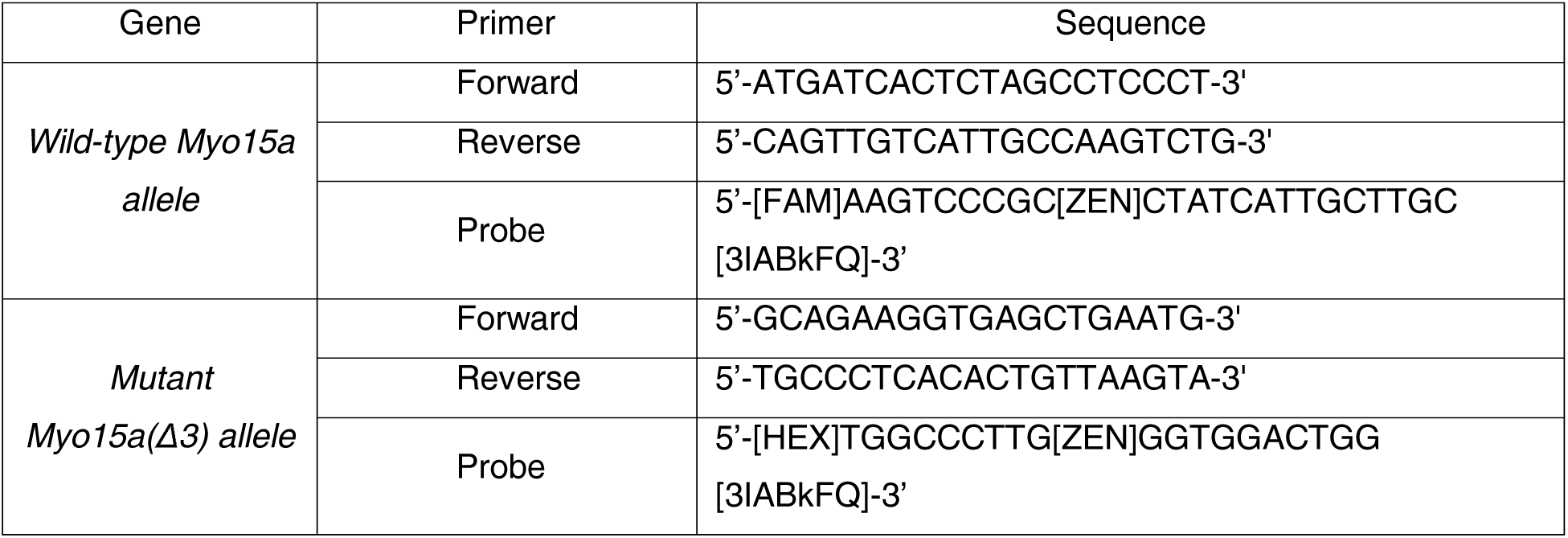
Primer Sequence to genotype *Myo15a-*Δ*3* allele.

### Quantitative polymerase chain reaction (qPCR)

Total RNA was extracted from C57BL/6J mice cochleae at postnatal days 0, 7, 14, 30, and 60. The otic capsule was quickly dissected in Leibovitz L-15 (Life Technologies) to isolate the cochlea and remove the vestibule and semi-circular canals, and snap-frozen in liquid nitrogen. RNA isolation was performed using TRI Reagent (Zymo Research), followed by phenol-chloroform separation and column purification using direct-zol RNA miniprep kit (Zymo Research), adapted from the protocol described (Vikhe Patil, Canlon, and Cederroth 2015). RNA quality was measured using a TapeStation (Agilent). Only samples with a RNA Integrity Number (RIN) value > 7 were used for analysis. First-strand cDNA was generated from total RNA using random-primer qScript cDNA synthesis (Quantabio). QPCR was performed using the SYBR Green dsDNA-binding dye to measure quantify DNA amplification. QCPR primers (Table 2) were designed using Primer3Plus and validated using BLAST searches against the mouse genome. For each reaction, first-strand cochlear cDNA was combined with a set of primers and PowerUp SYBR Green Mastermix (Applied Biosystems). PCR was performed on the QuantStudio 6 Real-Time PCR system (Applied Biosystems) using the following thermal profile: 50°C (2 min), 95 °C (2 min), followed by 40 cycles of 95 °C (15 s), 60 °C (1 min). Cycle thresholds (C_T_) for target genes were normalized (ΔC_T_) to hypoxanthine-guanine phosphoribosyltransferase (*Hprt*) as a housekeeping reference. ΔC_T_ values for each isoform were then normalized to P0 to yield ΔΔC_T_. Fold changes were calculated as 2^(-ΔΔC_T_). Three technical replicates were performed per target gene, and 3 independent biological determinations were analyzed per age group.

**Table 2.**
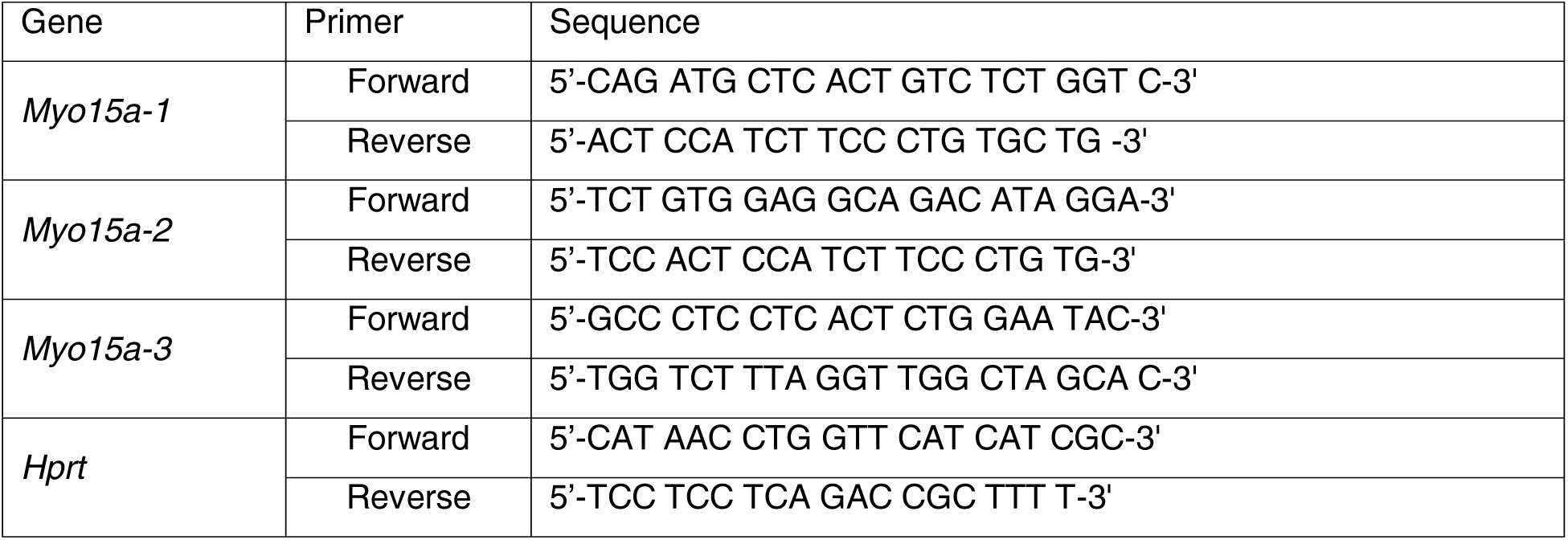
Primer sequences used for qPCR.

### BaseScope *in situ* hybridization

RNA *in situ* hybridization was conducted using the BaseScope RED assay protocol (ACD Bio) with modifications as described (Ghosh et al. 2022). Temporal bones were extracted, and cochleae perfused via the round window with 4% paraformaldehyde (PFA) (Electron Microscopy Sciences) for 3 hours at room temperature. Cochleae were micro-dissected in cold 20% RNAlater (Invitrogen), and isolated organ of Corti samples stored 96 well plates in RNAlater at 4 °C. For sample pretreatment, RNAlater was removed and replaced with 200 µL of 50% ethanol and incubated for 5 min at room temperature. Samples were taken through a graded ethanol series (70%, 95%, 100%) for 5 min each. Ethanol was then aspirated and pre-treatment I (hydrogen peroxide, ACD Bio) and incubated for 30 min at room temperature. Samples were then washed twice in MilliQ water for 3 mins each. After the final wash, samples were incubated with a 1:100 dilution of a citric acid-based antigen unmasking solution (H-3300, Vector Laboratories) in MilliQ water, for 5 min at 65 °C. Samples were washed extensively in MilliQ water. Following the last wash, samples were incubated with Protease Plus (ACD Bio) for 30 min at room temperature. Samples were washed with MilliQ water and stored at 4 °C overnight. The following day, probes were added to each sample and incubated at 40 °C in a humidified oven for 2 hours. Probes targeting specifically targeting each *Myo15a* isoform were designed by ACDbio; positive control (BA-Mm-Ppib-1ZZ), negative control (BA-DapB-1ZZ), *Myo15a-1* (BA-Mm-Myo15-tv1-E2E3), *Myo15a-2* (BA-Mm-Myo15-tv2-E1E2), *Myo15a-3* (BA-Mm-Myo15-E2aE3). After hybridization, tissues were washed 3 times with 1X wash buffer from the RNAscope wash buffer reagents (#320058, ACD Bio). Samples were then sequentially incubated with amplification reagents #1 - 8 from the BaseScope v2-RED (# 323910, ACD bio) at 40 °C for 30 mins. Samples were washed three times in 1X wash buffer between amplification steps. A RED working solution was prepared just before use by mixing BaseScope Fast RED-B and BaseScopeFast RED-A at a 1:60 ratio. Samples were incubated in RED solution for 10 min at room temperature, followed by 3 x 5 mins washes in MilliQ water. Samples were then incubated with 5 µg/mL of a rabbit polyclonal anti-MYO7A antibody (# 25-6790, PROTEUS) diluted in 0.1% Triton X-100 (Fisher Bioreagents) in PBS overnight at 4 °C. Following washes in PBS, samples were incubated with 2 µg/mL Alexa Fluor 488 Donkey anti-rabbit IgG (H+L) (# A-21206, Invitrogen) in PBS with 0.1% Triton X-100 for 2 hours at room temperature. After 3 times for 5 mins washes in PBS, samples were incubated with 1.5 µg/mL 4’,6-diamidino-2-phenylindole (DAPI) (#62247, Invitrogen) diluted in PBS for 15 min at room temperature. After 3 washes with PBS, tissues were mounted in Prolong Glass (#P36980, Invitrogen) and imaged using a confocal spinning disk microscope with a 60x oil immersion objective lens (Nikon, see description below). Z stacks were captured at 0.3 μm spacing.

### Auditory brainstem response (ABR)

ABR recordings were performed using a Tucker-Davis Technologies (Alachua, FL) system as previously described (Kim et al. 2023). Mice of both sexes were assembled into cohorts of *Myo15a^(+/+)^*, *Myo15a^(+/^*^Δ*3)*^ and *Myo15a^(^*^Δ*3/*Δ*3)*^ and longitudinally tested at P17, P30 and P60. Auditory measurements were performed blind to genotype. Mice were anesthetized through intraperitoneal injection with ketamine (100 mg/kg, Covetrus) and xylazine (10 mg/kg, Covetrus). Needle electrodes from the RA4PA-RA4LI system (Tucker-Davis Technologies) were placed sub-dermally at the vertex, ipsilateral (reference) and contralateral (ground) ears. A single MF1 speaker (Tucker-Davis Technologies) delivered auditory stimuli, with its output directed into the right ear canal via a connecting tube. A broadband impulse click stimulus, or pure-tone stimuli at 8, 16, and 32 kHz were delivered at different amplitudes to assess auditory brainstem responses. The ABR waveforms were recorded by averaging the electrical responses from electrodes over multiple stimulus presentations at each intensity level. The initial high-intensity level was set at 90 dB SPL and then decreased in 5 -10 dB increments until no response was detectable. The ABR threshold was defined as the lowest intensity level (in dB SPL) at which the peak in wave I could be detected. This process was repeated for each stimulus frequency to obtain frequency-specific ABR thresholds.

### Distortion product otoacoustic emissions (DPOAE)

DPOAE responses were measured using a Tucker-Davis Technologies (TDT) system according to the manufacturer’s DPOAE User Guide. The same cohort of mice utilized for longitudinal ABR measurements were also used for DPOAE testing at P40. Mice were anesthetized as described above. DPOAE stimuli were generated using SigGen software (TDT) and two MF1 speakers (TDT) connected to an ER10B+ probe (Etymotic Research) positioned in the right ear canal. The stimuli consisted of two pure-tone frequencies, designated f1 (the lower frequency) and f2 (the higher frequency), with an f2/f1 ratio of 1.2. These tones were geometrically centered around 8, 16, and 32 kHz. At each center frequency, the tone levels L1 and L2 remained equal and were reduced in 10□dB steps from 80 to 20□dB SPL. Stimuli ranging from 20-80 dB SPL were presented, whilst recording the distortion product generated at 2f1-f2. DPOAE thresholds were defined as the lowest level at which the 2f1-f2 distortion product was at least 5 dB above the surrounding noise floor (Cederholm et al. 2012). The noise floor was calculated as the mean amplitude across at least ten neighboring frequencies to ensure a robust comparison base (Powers et al. 2006).

### Hair cell injectoporation

Cochlear hair cells were injectoporated with plasmid DNA as described (Xiong et al. 2014). Briefly, the sensory epithelium was dissected from C57BL/6J mice at P5 in Hank’s balanced salt solution (#14025092, Life Technologies), and the cochlear duct was opened by making an incision between Reissner’s membrane and the stria vascularis. The tissue was unfolded allowing the cochlear lateral wall to adhere to a tissue-culture treated polystyrene dish (#CC7672-3359, USA Scientific) containing DMEM/F12 (Thermo Fisher Scientific, Cat. # 11039047) with 0.1 mg/mL penicillin. The culture was incubated at 37 °C with 5% CO_2_ for 2 hours before injectoporation was performed. A glass micropipette with a 2 µm tip diameter loaded with plasmid DNA (2 mg/mL in water) was oriented perpendicular to the IHC row. The tip of the micropipette was inserted into the space between two IHCs, and pressure was supplied by a microinjector to inject plasmid into the tissue. An electroporator (ECM 830, BTX) delivered a series of three 60V square-wave electrical pulses (15 ms) at 1 second intervals to platinum wire electrodes spaced 2 mm apart and positioned directly across the microinjection site. After the electroporation, the culture media was exchanged with Neurobasal-A (#12349015, Life Technologies) supplemented with 2 mM L-glutamine (#25030081, Life Technologies), 1x N2 supplement (#17502048, Life Technologies), 75 mg/mL D-glucose (# 410955000, Life Technologies), and 0.1 mg/mL penicillin. Cultures were cultured for 18 h at 37°C in 5% CO_2_ incubator, then fixed with 4% paraformaldehyde (#15710, Electron Microscopy Sciences) in PBS for 2 hours. Fixed samples were stained with Alexa Fluor 568 phalloidin (0.5 U/mL, A12380, Life Technologies) in PBS with 0.1% Triton X-100 (X-100-100ML, Sigma) at room temperature for 1 h. The tectorial membrane was removed, and the tissue was mounted in Prolong Diamond (P36961, Life Technologies). After curing, the slides were imaged with a Leica Plan Apo 63x/1.40 NA oil immersion objective on Leica SP8 inverted confocal microscope (Leica Microsystems) operating in resonant scanning mode and deconvolved using Leica LIGHTNING deconvolution with the default settings.

### Whole mount immunofluorescence

Cochlear immunofluorescence was performed as previously described (Fang et al. 2015). Temporal bones were micro-dissected in Leibovitz L-15 media (10-045-CV, Corning) to expose the cochlea and transferred to 4% paraformaldehyde in PBS for 30 min. A small hole was punched at the helicotrema, and fixative was gently perfused through the oval and round windows using a 200 µL pipette to ensure rapid fixation. After washing in PBS, cochleae were micro-dissected to remove the stria vascularis, Reissner’s and tectorial membranes. Samples were permeabilized in PBS with 0.5% Triton X-100 for 30 min and blocked for 1 hour in PBS with 2% bovine serum albumin (BP1605-100, Fisher Scientific). Samples were incubated overnight at 4 °C with primary antibodies diluted in blocking solution. Primary antibodies were as follows: PB48 rabbit anti-MYO15A (Liang et al. 1999), PB888 rabbit anti-MYO15A-1 (Fang et al. 2015), TF1 rabbit anti-MYO15A (Liang et al. 1999), HL5136 rabbit anti-WHRN (Belyantseva et al. 2005), mouse anti-EPS8 (610143, BD Biosciences). After extensive washing in PBS, samples were incubated with 1 μg/mL of either Alexa Fluor 488 donkey anti-rabbit IgG (A-21206, Life Technologies) or Alexa Fluor 488 donkey anti-mouse IgG (A-21202) secondary antibodies for 2 hours at room temperature. Samples were washed 3 x 5 mins in PBS and incubated a 1:200 dilution of phalloidin CF 647 (00041, Biotium) in PBS for 30 mins. Following washing in PBS, samples were mounted under high precision #1.5 cover glass (Thorlabs) using ProLong Glass (P36980, Life Technologies). Samples were visualized using an inverted microscope (Nikon Ti2-E) equipped with 100x oil objective (CFI Apochromat TIRF, 1.49 N.A., Nikon), spinning disk confocal unit (CSU-X1, Yokogawa) with a super-resolution Live-SR module (Gataca). Images were captured using a Prime 95B (Teledyne Photometrics), or Quest II (Hamamatsu) controlled by NIS-Elements (AR version 5.2, Nikon). Z-stacks were captured with a 0.1 μm spacing. For quantification of EPS8, WHRN or MYO15A immunofluorescence, up to 10 circular regions of interest (ROI; 0.4 µm^2^) were placed at the tips of row 1 stereocilia in each hair cell. The Z-stack position was adjusted for each ROI to obtain the maximum fluorescence signal, and the mean row 1 intensity was calculated per hair cell. Comparisons between different genotypes were made using identical fluorophores, laser excitation and camera exposure, and absolute fluorescence values were normalized relative to *Myo15a^(+/^*^Δ*3)*^ samples. For measurements of PB888 fluorescence intensity, the ratio of row 1 / row 2 was calculated for each sample and not normalized.

### Mammalian cell culture and transfection

The pEGFP-C2-Myo15a-2 plasmid was a gift from Thomas Friedman and expresses a fusion of EGFP with mouse MYO15A isoform 2 (NP_874357.2) (Belyantseva, Boger, and Friedman 2003). The pEGFP-C-Myo10-HMM plasmid was a gift from Thomas Friedman and expresses a fusion of EGFP with a truncated mouse MYO10 (NP_062345) (Bird et al. 2017). The pmCherry-C1-EPS8 plasmid was a gift from Christien Merrifield (Addgene plasmid #29779). All expression plasmids were verified by Sanger sequencing (Eurofins Genomics) and endotoxin free plasmid DNA prepared using a ZymoPure II plasmid midiprep kit (D4200, Zymo Research). HeLa cells (CCL-2) were obtained from the ATCC and cultured in Dulbecco’s Modified Eagle Medium (DMEM) (11965118, Life Technologies) supplemented with 10% heat-inactivated fetal bovine serum (FBS) (Sigma-Aldrich) and 1% GlutaMax supplement (35050079, Life Technologies). Cells were maintained at 37°C in a humidified 5% CO_2_ atmosphere. HeLa cells were transfected at 70-80% confluency using polyethyleneimine (PEI Max, 24765-1, PolySciences Inc). The transfection mixture was prepared in a serum-free DMEM with a 4:1 molar ratio of PEI to DNA, incubated for 20 mins at RT, and added to cells drop-wise. After incubation overnight, transfected HeLa cells were trypsinized and seeded onto glass-bottom dishes (D35-20-1.5-N, Cellvis) coated with 10 µg/mL human fibronectin (F0895, Sigma-Aldrich). Cells were allowed to adhere for 5 hours before imaging live on an inverted microscope (Nikon Ti2-E) equipped with a 60x oil immersion objective (CFI Plan Apo Lambda 60x, 1.4 N.A) and a confocal spinning disk microscope, as described above.

### Analysis of fluorescence colocalization in filopodia

Line scans were traced along individual filopodia based on the fluorescence signal of the phalloidin channel. This process was carefully conducted with the hidden fluorescence of the bait (Myosin) and prey (EPS8/WHRN) to eliminate biased filopodia selection. Only filopodia extending more than 2 µm from the cell body were considered to ensure meaningful comparisons between the tips and shafts of the filopodia. 10 filopodia per cell were imposed with a comprehensive analysis encompassing 300 filopodia per interaction derived from three independent experiments. Data from the bait and prey fluorescence line scans were exported to a tab-delimited text file for further analysis. As previously described, a custom-built MATLAB tool with a graphical user interface was used to measure fluorescence signal correlation along the filopodia shaft (Bird et al. 2017). The interaction index between bait (myosins) and prey (EC) is assessed using a spatial correlation analysis algorithm. Pearson’s correlation coefficient (r) is computed along the filopodia shaft to gauge the relationship between the fluorescence intensities of myosins and EC. The probability that the observed correlation occurs by random chance is determined via bootstrapping, which randomizes the EC signal. A filopodium is considered to exhibit a positive myosin-EC interaction if: 1. the p-value is less than 0.01, and 2. the prey intensity at the filopodial tip surpasses the shaft signal by 6 standard deviations (SDs). The interaction index is then calculated as the ratio of filopodia with positive interactions to the total number of filopodia measured.

### Scanning electron microscopy (SEM)

Samples for SEM were prepared and imaged as previously described (Vélez□Ortega et al. 2017). Briefly, organ of Corti explants were fixed overnight at 4 °C in a fixative containing 3% paraformaldehyde and 5% glutaraldehyde in 0.1 M sodium cacodylate buffer (PH 7.4, Electron Microscopy Sciences) supplemented with 2 mM CaCl_2_. Explants were then dehydrated with a graded series of ethanol, critical point dried from liquid CO_2_ (Leica EM CPD300), followed by sputter coating with 5 nm of platinum controlled by a thin-film thickness monitor (Q150T, Quorum Technologies, Guelph, Canada). A field-emission scanning electron microscope (Helios Nanolab 660, FEI, Hillsboro, OR) was used to image the hair bundles. Quantitative measurements of stereocilia diameter were performed using ImageJ (version 2.14) as previously described (Vélez□Ortega et al. 2017).

### Focused ion-beam scanning electron microscopy (FIB-SEM)

Temporal bones were dissected in chilled Leibovitz (L-15) medium immediately following euthanasia. The cochleae were perfused through the oval window using a fixative solution of 3% paraformaldehyde (PFA) and 3% glutaraldehyde (GLU) in 0.1 M sodium cacodylate buffer (pH 7.4) and 1% tannic acid for 24 h at 4 °C. Cochleae were stored in a 1:20 diluted solution of distilled water to 3% PFA/GLU without tannic acid. Organ of Corti explants were dissected and cryoprotected with a glycerol series (5-25%) before overnight incubation in 30% glycerol at room temperature. The tissue was then plunge frozen in liquid N_2_, freeze-substituted in methanol with uranium acetate, and low temperature embedded in Lowicryl HM20. The embedded resin blocks were mounted on low profile 45/90° SEM pins, then trimmed on an RMC PowerTome XL microtome and sputter-coated with 25 nm platinum (Q150T, Quorum Technologies, Guelph, Canada). The samples were imaged with a field-emission scanning electron microscope (SEM) (Helios Nanolab 660, FEI, Hillsboro, OR) equipped with a focused ion beam (FIB). Serial sections of 20 nm were milled from the block face while alternating imaging with the scanning electron beam with a ∼1.4 nm/pixel resolution.

### Image processing and statistical analysis

Z-stack images were processed for deconvolution and denoising using a blind algorithm within the NIS-Elements Advanced Research software. Subsequent image analysis and processing were performed with Prism (GraphPad) and ImageJ software (https://imagej.nih.gov). To compare two groups with one variable, an unpaired two-tailed t-test was used if the data were normally distributed, and a Mann–Whitney U test was used if the data were not normally distributed. For comparisons of more than two samples with one variable, a one-way ANOVA followed by Dunnett’s, Šidák, Fisher’s LSD, or Dunn’s multiple comparison tests was conducted. For comparisons involving two variables, a two-way ANOVA followed by Tukey’s, Šidák, or Fisher’s LSD multiple comparison tests was utilized. Quantitative data are presented as mean ± SD. Statistical significance was defined as a p-value less than 0.05, indicated as follows: *<0.05, **<0.01, ***<0.001, and ****<0.0001.

## DATA AVAILABILITY

The raw data that support the findings of this study are available from the corresponding authors upon request.

## ACKNOWLEDGEMENTS

We thank Randall Harley for cloning the pEGFP-C-MYO15A-3 plasmid. We are also grateful to Peter Barr-Gillespie and members of the Bird Lab for critical reading of the manuscript and insightful discussions. This research was supported by National Institutes of Health awards R01 DC 018827 (JEB). The content is solely the responsibility of the authors and does not necessarily represent the official views of the National Institutes of Health. The funding body had no role in study design, data collection or analysis, decision to publish, or preparation of the manuscript.

## AUTHOR CONTRIBUTIONS

Conceptualization: JEB, GIF

Investigation: GB, AD, SH, XL, MJK

Formal analysis: GB, AD, XL, MJK

Writing - original draft: GB, JEB

Writing – review and editing: All authors.

Project administration: JEB

Funding acquisition: JEB

---

**Figure S1.**
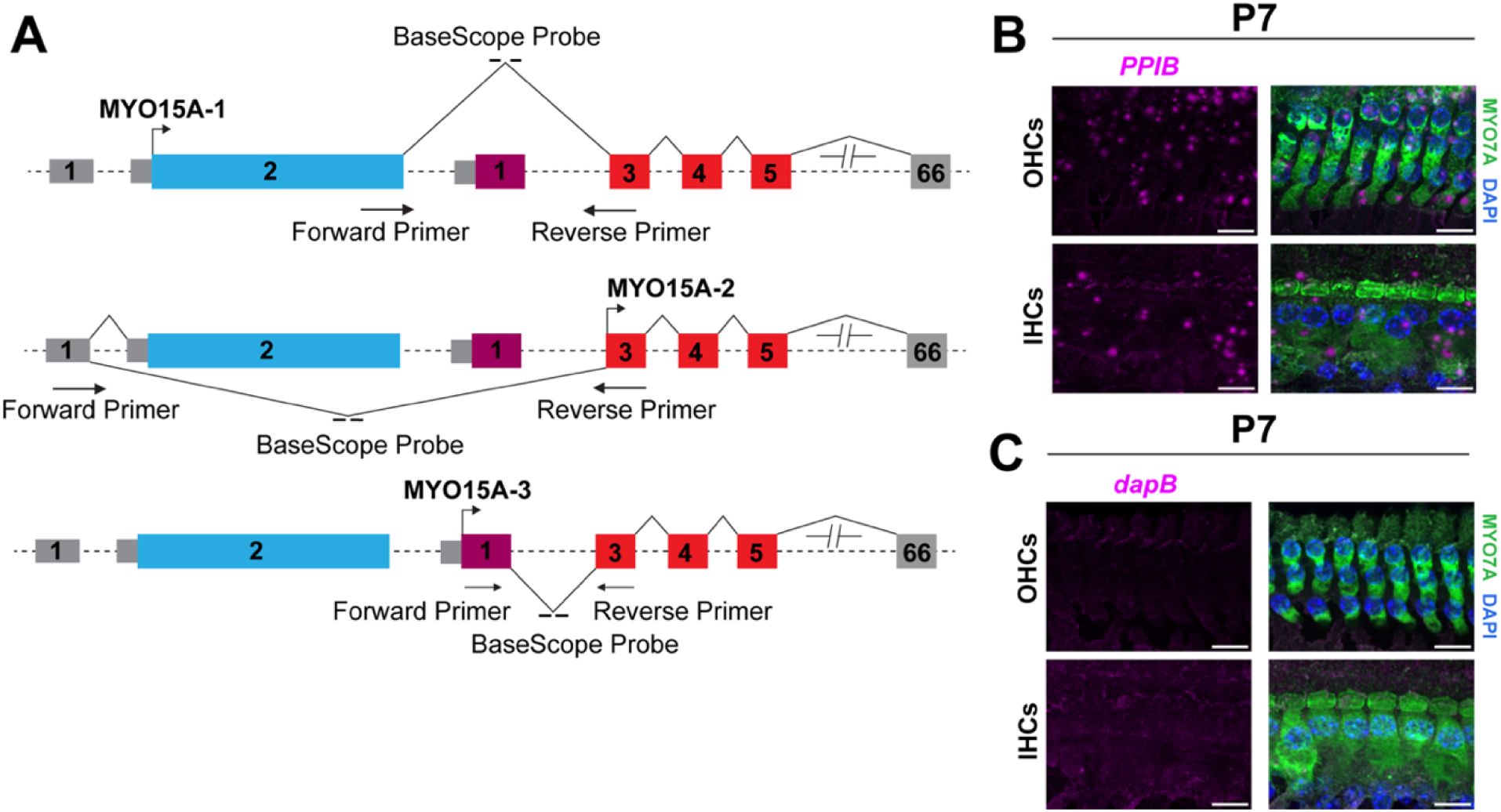
*Myo15a* isoforms are expressed at distinct stages during cochlear development. **(A)** Schematic representation of *Myo15a* gene structure with alternative transcription start site. Forward and reverse primers were designed to specifically detect splicing between exon 2 and 3 (*Myo15a-1*), exon1 and 3 (*Myo15a-2*), and to amplify the region spanning exons 1 and 3 (*Myo15a-3*). BaseScope probes were designed to span the exon junctions. **(B)** *PPIB* (positive control) and **(C)** *DAPB* (negative control) BaseScope signal to validate probe specificity.

**Figure S2.**
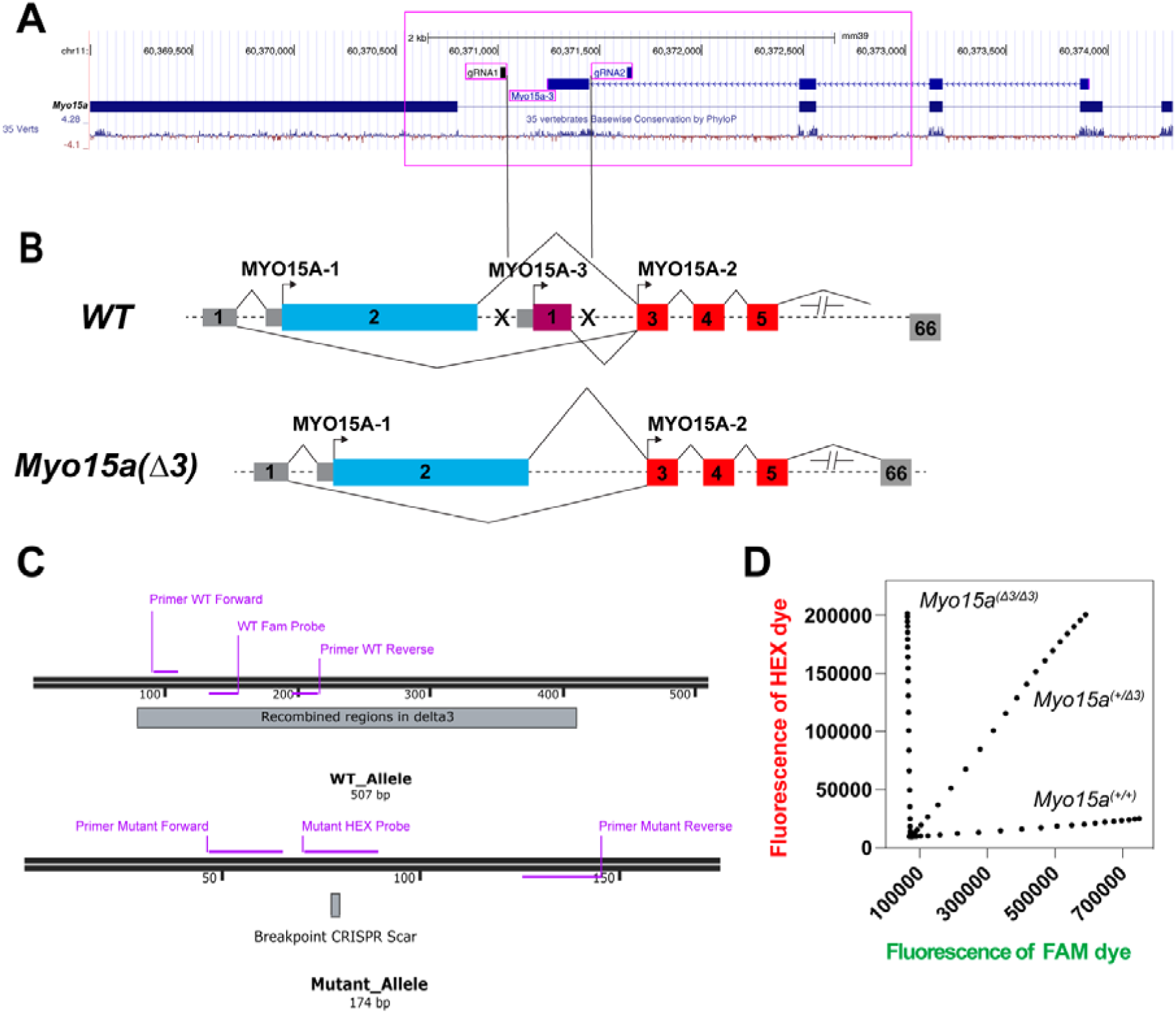
Generation and genotyping of *Myo15a-*Δ*3* mice. **(A)** Positions of two guide RNAs (gRNAs) on mouse genome targeting the flanking regions of the exon1 of *Myo15a-3*. **(B)** Schematic representation of the *Myo15a* gene locus. The unique exon 1 of the *Myo15a-3* was excised using CRISPR/Cas9, resulting in the *Myo15a^(^*^Δ*3/*Δ*3)*^ mutant allele. **(C)** Genotyping strategy for *Myo15a^(^*^Δ*3/*Δ*3)*^ mice. A multiplex TaqMan assay was used to simultaneously detect the wild-type (WT) and mutant alleles in a single reaction. **(D)** The probes were labeled with distinct fluorophores: 6-FAM (fluorescein, FAM) for the WT allele and hexachlorofluorescein (HEX) for the mutant allele, allowing for discrimination between genotypes.

**Figure S3.**
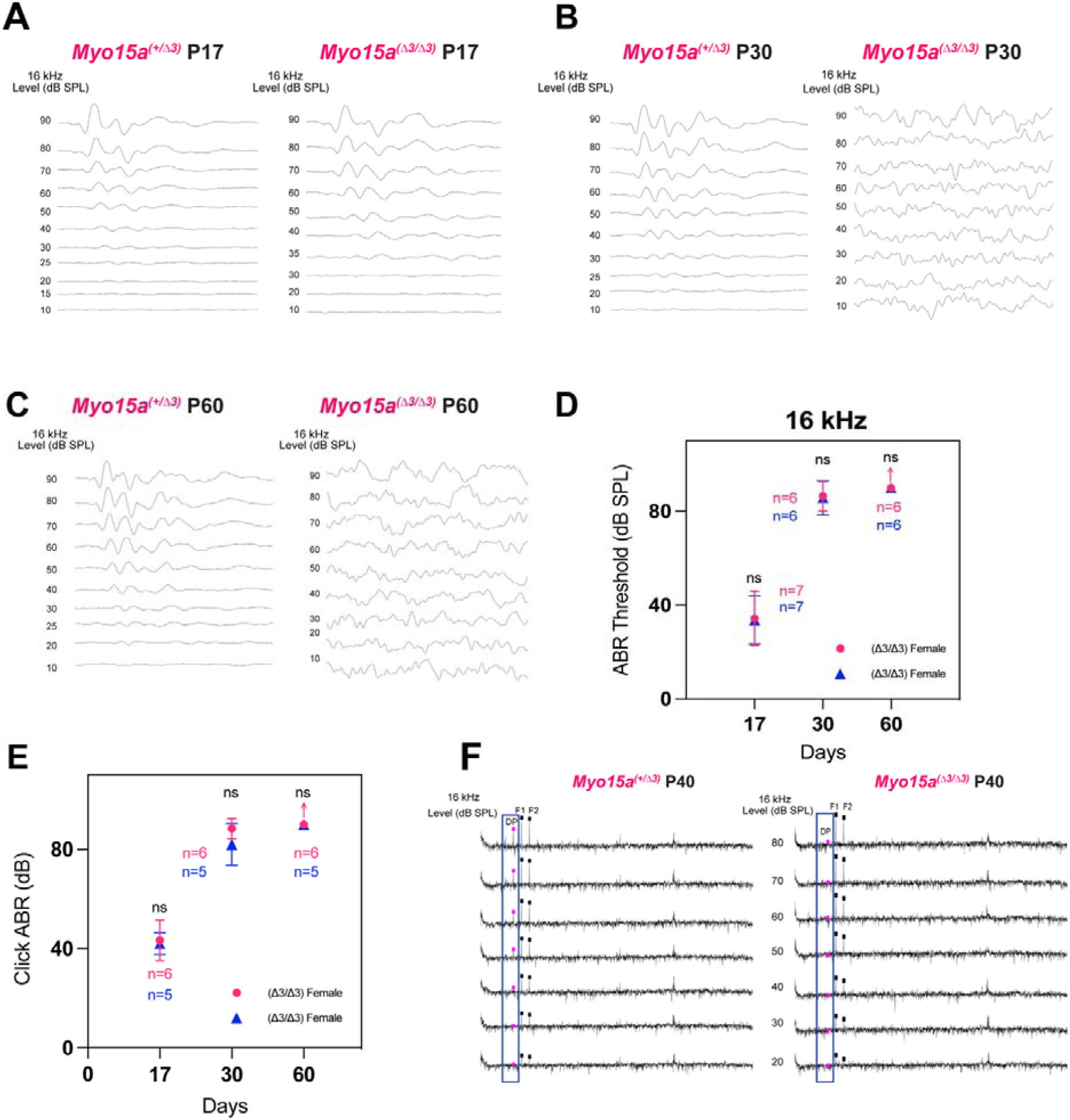
**(A-C)** Comparison of ABR waveforms at 16kHz between *Myo15a^(+/^*^Δ*3)*^ and *Myo15a^(^*^Δ*3/*Δ*3)*^ across ages. **(D, E)** Comparison of ABR thresholds at 16kHz and click ABR between male and female of *Myo15a^(^*^Δ*3/*Δ*3)*^ mice. **(F)** DPOAE gram illustrates the frequency response of the distortion product emissions. Frequencies f1 and f2 are labeled, and the resulting distortion product (DP) is shown for both genotypes.

